# Enhanced mTORC1 signaling and Protein Synthesis in Parkinson’s Disease Pathogenesis Disease Pathogenesis

**DOI:** 10.1101/2022.10.03.510455

**Authors:** Mohammed Repon Khan, Xiling Yin, Sung-Ung Kang, Jaba Mitra, Hu Wang, Saurav Brahmachari, Senthilkumar S. Karuppagounder, Yasuyoshi Kimura, Aanishaa Jhaldiyal, Hyun Hee Kim, Hao Gu, Rong Chen, Javier Redding-Ochoa, Juan Troncoso, Taekjip Ha, Valina L. Dawson, Ted M. Dawson

## Abstract

Pathologic α-syn destabilizes the TSC 1 and 2 complex leading to mTORC1 activation, enhanced protein translation and neurodegeneration in PD.

**Abstract:** Pathological α-synuclein (α-syn) plays an important role in the pathogenesis of α-synucleinopathies such as Parkinson’s disease (PD). Disruption of protein homeostasis is thought be central to PD pathogenesis, however the molecular mechanism of this deregulation is poorly understood. Here we report that pathologic α-syn binds to tuberous sclerosis protein (TSC) 2 and destabilizes the TSC1-TSC2 complex leading to activation of the mechanistic target of rapamycin (mTOR) complex 1 (mTORC1) and enhanced mRNA translation. Dopamine neuron loss, behavioral deficits and aberrant biochemical signaling in the α-syn preformed fibril (PFF) and *Drosophila* α-syn transgenic models of pathologic α-syn induced degeneration were attenuated by genetic and pharmacologic inhibition of mTOR and protein translation. Our findings establish a potential molecular mechanism by which pathologic α-syn activates mTORC1 leading to enhanced protein translation and concomitant neurodegeneration in PD.

## INTRODUCTION

Parkinson’s disease (PD) is a major neurodegenerative disorder that is increasing in prevalence and incidence (*1*). PD is characterized by the loss of dopamine (DA) neurons leading to the major motor symptoms including rest tremor, rigidity, bradykinesia and postural instability (*2*). In addition to loss of DA neurons, there is widespread deficits in other neuronal systems that account for the non-motor features of PD (*3*).

Central to the disease progression is the accumulation and the formation of pathologic α-synuclein (α-syn). How pathologic α-syn leads to the degeneration in PD is poorly understood. Pathologic α-syn can lead to defects in protein homeostasis (*4*). A molecular mechanism accounting for this dysregulation is currently lacking. Defects in mRNA translation are linked to autosomal dominant PD (*5, 6*) and autosomal recessive PD (*7*), where there is aberrant bulk protein synthesis that is coupled to neurodegeneration (*8, 9*). This aberrant protein synthesis is, in part, mediated by dysregulation of LRRK2 phosphorylation of ribosomal proteins (*5, 6*) and PTEN-induced kinase 1 (PINK1) and parkin modulation of eukaryotic initiation factor 4E (eIF4E)-binding protein (4E-BP) (*7*). Whether aberrant protein translation plays a role in role in α-syn pathogenesis is not known. Here we establish a connection between pathologic α-syn and deregulation of mRNA translation in PD and reveal a molecular mechanism in which pathologic α-syn directly binds tuberous sclerosis protein 2 (TSC2) and destabilizes the TSC1 and TSC2 complex. This destabilization of the TSC1-TSC2 complex, triggers aberrant mammalian target of rapamycin (mTOR) complex 1 (mTORC1) activation, which results in enhanced mRNA translation that contributes to pathologic α-syn induced neurodegeneration.

## RESULTS

### Pathologic α-syn Biochemically Interacts with the Critical Components of Translation and mTOR Regulatory Pathways

Proteins interacting with pathologic α-syn were evaluated using two complementary proteomic approaches. First, overexpression of human A53T α-syn was tagged with APEX (human A53T α-syn-APEX) and used to examine human A53T α-syn interacting proteins in HEK293 cells via mass spectrometry (Figure 1A). Second, biotin labeled human α-syn preformed fibrils (PFF) and non-labeled human α-syn PFF were stereotactically injected into mouse striatum and subjected to streptavidin immunoprecipitation followed by mass spectrometry identification (Figure 1B). In HEK293 cells expressing human A53T α-syn-APEX, 2015 proteins were identified in the presence of hydrogen peroxide (H_2_O_2_) after subtracting interacting proteins that were detected in the absence of H_2_O_2_ (Figure 1C and Data Set S1). In mouse striatum, 932 α-syn PFF interacting proteins were identified after subtracting the proteins identified from non-labeled human α-syn PFF from the biotin labeled human α-syn PFF (Figure 1C and Data Set S2). In the mass spectrometric analysis, only proteins that were at identified via at least 2 separate peptides of at least 6 amino acids and enriched at least 2-fold were included as positive interactors in the human A53T α-syn-APEX (Figure 1C and Data Set S1) and the α-syn PFF pull downs (Figure 1C and Data Set S2). These interacting proteins from both approaches were subjected to major pathways and Gene Ontology (GO) analysis and in both data sets RNA processing and translation initiation were the highest biologic processes that were identified (Table S1 and Table S2). 256 proteins were found to be common in both data sets (Figure 1C). GSA (Gene set analysis) in Cytoscape was used for pathway analysis using the 256 common interacting proteins (Figure 1D). Translation pathway and mTOR pathways were found to be statistically enriched in GSA pathway analysis (Figure 1D). Immunoblot analysis of streptavidin IP from human A53T α-syn-APEX peroxidase labelling and α-syn-PFF pulldown from mouse striatal tissues was able to confirm that mTOR pathway protein partners are present in both approaches (Figures 1E and 1F). Regulation of catabolic processes that included autophagy were also statistically enriched in the GSA pathway analysis (Figure 1D) consistent with the role of endolysosomal pathways and disruption of macroautophagy and chaperone mediated autophagy in the pathogenesis of pathologic α-syn (*10, 11*). Taken together, these pulldown and proteomics data suggest that pathologic α-syn biochemically interacts with the components of translation and mTOR regulatory pathways. Moreover, the proteomic analysis confirms the importance of autophagy in the pathogenesis of pathologic α-syn. Since little is known about how pathologic α-syn might regulate mRNA translation and mTOR pathways a series of experiments were conducted to explore the effect of pathologic α-syn on mRNA translation and mTOR pathways.

**Fig. 1:**
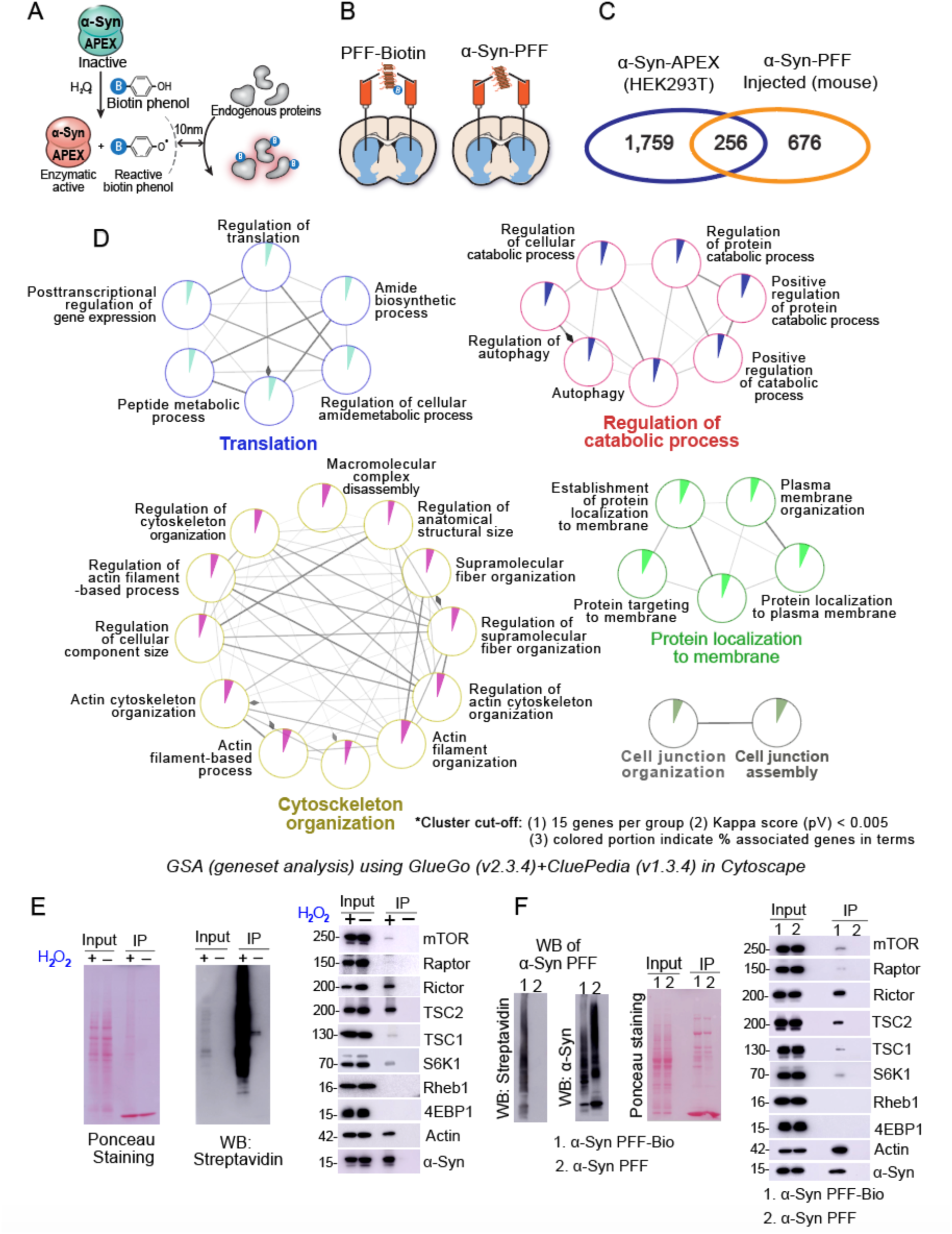
Pathogenic α-syn biochemically interacts with the critical components of translation and mTOR pathways. (**A**) Schematic of proteomics assay design for human A53T α-syn-APEX (Ascorbate Peroxidase) proximity labelling of interacting proteins. (**B**) Schematic of stereotactic injection of human α-syn PFF into mouse striatum. (**C**) Venn-diagram shows unique and common interacting protein partners. (**D**) GSA (Gene Set Analysis) in Cytoscape shows the major pathways that emerged from common interacting proteins, which include translation, regulation of catabolic process, cytoskeletal organization, and cell junction assembly. (**E**) α-syn directly interacts with critical components of mTOR pathway. Left two panels show input and immunoprecipitation of biotinylated protein from HEK293 cell lysates. Right panel shows immunoblots of input and interacting proteins in streptavidin IP. (**F**) Immunoblots of biotinylated and non-biotinylated human α-syn PFF injected into mouse striatum. Ponceau staining shows streptavidin IP and input. The immunoblot panel at the right shows pulldown of critical component of mTOR pathways.

### Pathologic α-syn increases while monomeric α-syn represses protein translation

First, cell free in vitro and in vivo translation assays were performed to examine the effect of pathologic α-syn on protein translation and monomeric α-syn was used as a control. In a cell free in vitro translation (IVT) assay, both WT α-syn PFF and A53T α-syn PFF increased protein translation in a concentration dependent manner (Figures 2A-2C). Translation in vivo can be monitored in *Drosophila* by feeding radioactive S35 (*5*). De novo protein synthesis was quantified in UAS α-syn transgenic flies using both TH (TH Gal4> WT α-syn; TH Gal4> A53T α-syn) and pan-neuronal Elav (Elav Gal4> WT α-syn; Elav Gal4> A53T α-syn) drivers by S35 radioactive pulse labelling in fly brain tissues (Figure 2D and 2E). Both WT α-syn and A53T α-syn expression caused a significant increase in protein translation in both dopaminergic and pan-neuronally expressed α-syn flies. Transgenic fly brains were directly autoradiographed after S35 feeding providing a confirmation of the increase in protein translation (Figure 2F and 2G). Polysome profiles provided an additional confirmation of increased mRNA translation in transgenic fly brain homogenates as revealed by both WTα-syn and A53T α-syn brains showing increased abundance of polysome associated RNA in brain lysates (Figure 2H). The effect of α-syn PFF on translation was also examined in α-syn PFF treated primary cortical neurons using the SUnSET assay (*12*) (Figure 2I). Both puromycin immunoblot and immunohistochemistry showed a significant increase in translation (Figures 2I and 2J). The increase in neuronal translation was α-syn PFF concentration dependent (Figures 2K). To examine the effect of endogenous pathologic α-syn on protein translation, human A53T pathologic α-syn was isolated from late symptomatic human A53T α-syn transgenic brainstem (*13*) and added to primary neurons and subjected to the SUnSET assay (Figures 2L). Similar to the effect of recombinant pathologic α-syn in the IVT assay, endogenous human A53T pathologic α-syn showed a significant increase in mRNA translation (Figures 2L). While pathologic α-syn enhanced mRNA translation, exogenous recombinant human WT and A53T α-syn monomeric proteins repressed protein translation in the IVT assay in a concentration dependent manner (Figure S1A). Using a cell based puromycin SUnSET assay to monitor protein translation in cells (*12*), overexpression of WT and A53T α-syn in HEK293 cells also caused a significant repression of protein translation (Figure S1B). Polysome profiling of WT and A53T α-syn transfected HEK cells lysates revealed less RNA association with polyribosomes compared to the GFP transfection, which is consistent with reduced protein translation (Figure S1C). The repression of protein translation by α-syn monomeric proteins is consistent with a prior report (*14*). Taken together, these in vitro and in vivo translation data are supportive of pathologic α-syn increasing protein translation while monomeric human WT α-syn and A53T α-syn represses protein translation.

**Fig. 2:**
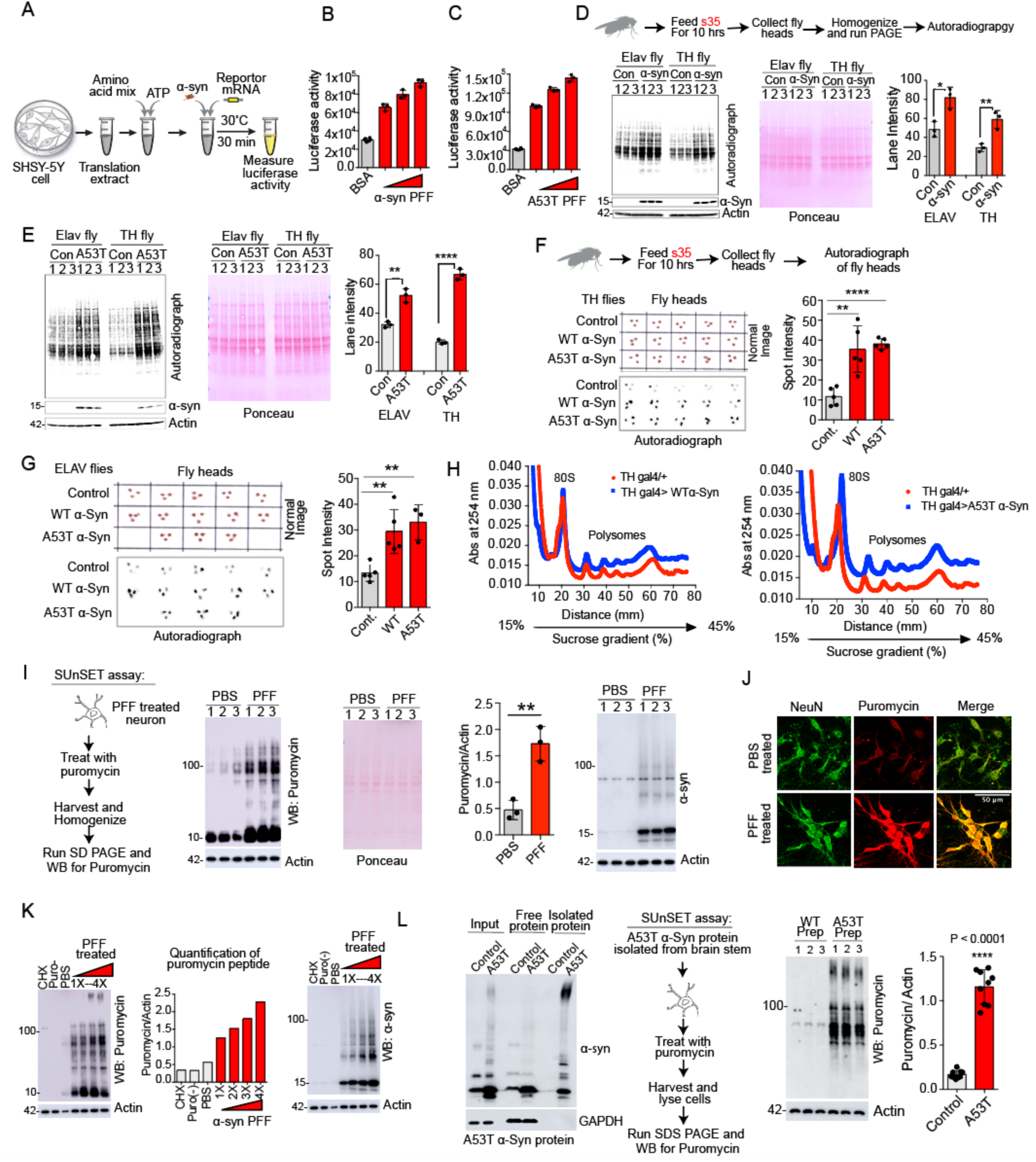
Pathogenic α-syn enhances mRNA translation. (**A**) Schematic diagram of cell free In Vitro Translation (IVT) assay design. (**B&C**) Quantification of luciferase activity in IVT assay by expressing cap dependent firefly luciferase reporter mRNA in presence of exogenously added WT and A53T α-syn PFF to IVTs (n=3). **(D)** Schematic shows radioactive S35 assay using TH Gal4>WT α-syn and Elav Gal4>WT α-syn flies. Autoradiograph shows incorporation of S35 in newly synthesized protein in fly brain tissues and ponceau staining shows protein loading. Graph shows quantification of newly synthesized protein (n=3). (**E**) Autoradiograph shows S35 incorporation in A53T α-syn brains. Ponceau staining shows protein loading and bar graph shows quantification of newly synthesized protein (n=3). (**F**) Schematic shows feeding of radioactive (S35) amino acid and autoradiography of intact fly brains using TH Gal4 expression system. Normal image and autoradiograph show S35 signal in fly brain and bar graph shows quantification of S35 incorporation (n=5). (**G**) Normal image and autoradiographs of fly brains under Elav Gal4 expression system. Bar graph shows quantification (n=3). (**H**) Polysome profiles show association of mRNA with polysome in fly brain lysates. (**I**) Schematic depiction of SUnSET assay design using α-syn PFF primary neurons. The immunoblot was probed for puromycin and α-syn. The bar graph shows quantification puromycin peptides (n=3). (**J**) Puromycin immunostaining of α-syn PFF treated primary cortical neurons. (**K**) Pathogenic α-syn mediates translation activation in concentration dependent manner in SUnSET assay in neuron. The immunoblot was probed for puromycin and α-syn. Bar graph shows quantification of normalized puromycin incorporation in neuron. (**L**) Left immunoblot shows detection of endogenous α-syn isolated from the brain stem of late symptomatic A53T transgenic mice. GAPDH used as a loading control. Schematic shows SUnSET assay design with isolated α-syn in primary neurons. Puromycin immunoblot shows newly synthesized puromycin peptides. The bar graph shows quantification of puromycin peptides in the western blot (n=9). For the figures 2D, 2E, 2I and 2L statistical significance was measured by unpaired two-tailed t-test. Data are expressed as mean ± SEM; and *p ≤ 0.05, **p ≤ 0.01, and ***p ≤ 0.001. For the figures 2F and 2G statistical significance was measured by One-way ANOVA, data are expressed as mean ± SEM, *p < 0.05, **p < 0.01, ****p < 0.0001.

### Pathologic α-syn Increases Protein Translation via mTOR Kinase

Second, experiments were performed to evaluate whether the increase in translation is dependent on mTOR kinase activity. Treatment of α-syn flies (TH Gal4>WT α-syn) with mTOR inhibitor, rapamycin diminishes the increase in protein translation to control fly levels (Figures 3A and 3B). The global translation inhibitor, anisomycin also reduced protein translation in both TH Gal4>WT α-syn and control flies (Figures 3A and 3B). These data led to the examination of the phosphorylation status of mTORC1 downstream substrates in the cell free IVT assays, in cellular and in vivo models of pathologic α-syn induced toxicity and in human PD postmortem brains. Treatment of IVT reactions with WT α-syn PFF and A53T α-syn PFF showed a significant increase in phosphorylation of mTOR on S2481 and the mTORC1 effectors ribosomal S6 kinase p70 (S6K) on T389 and eukaryotic translation initiation factor 4E-binding protein 1 (4E-BP1) and T37/46) (Figures S2A and S2B). Likewise, there was a significant increase in phosphorylation of mTORC1 effectors in α-syn PFF treated mouse primary cortical neurons (Figure S2C) and in the brain stem region of late symptomatic A53T α-syn transgenic mice (Figure S2D), while we were not able to detect mTORC1 pathway activation in brain stem from pre-symptomatic A53T α-syn mice (*13*) (Figure S2E). Consistent with the possibility that mTOR is activated in human PD, a significant increase in 4E-BP1 phosphorylation was observed in postmortem PD cingulate cortex, a region with substantial pathologic α-syn (*15*) as evidenced by elevated phospho serine (P-S)129 α-syn compared to control brains (Figure S2F).

**Fig. 3:**
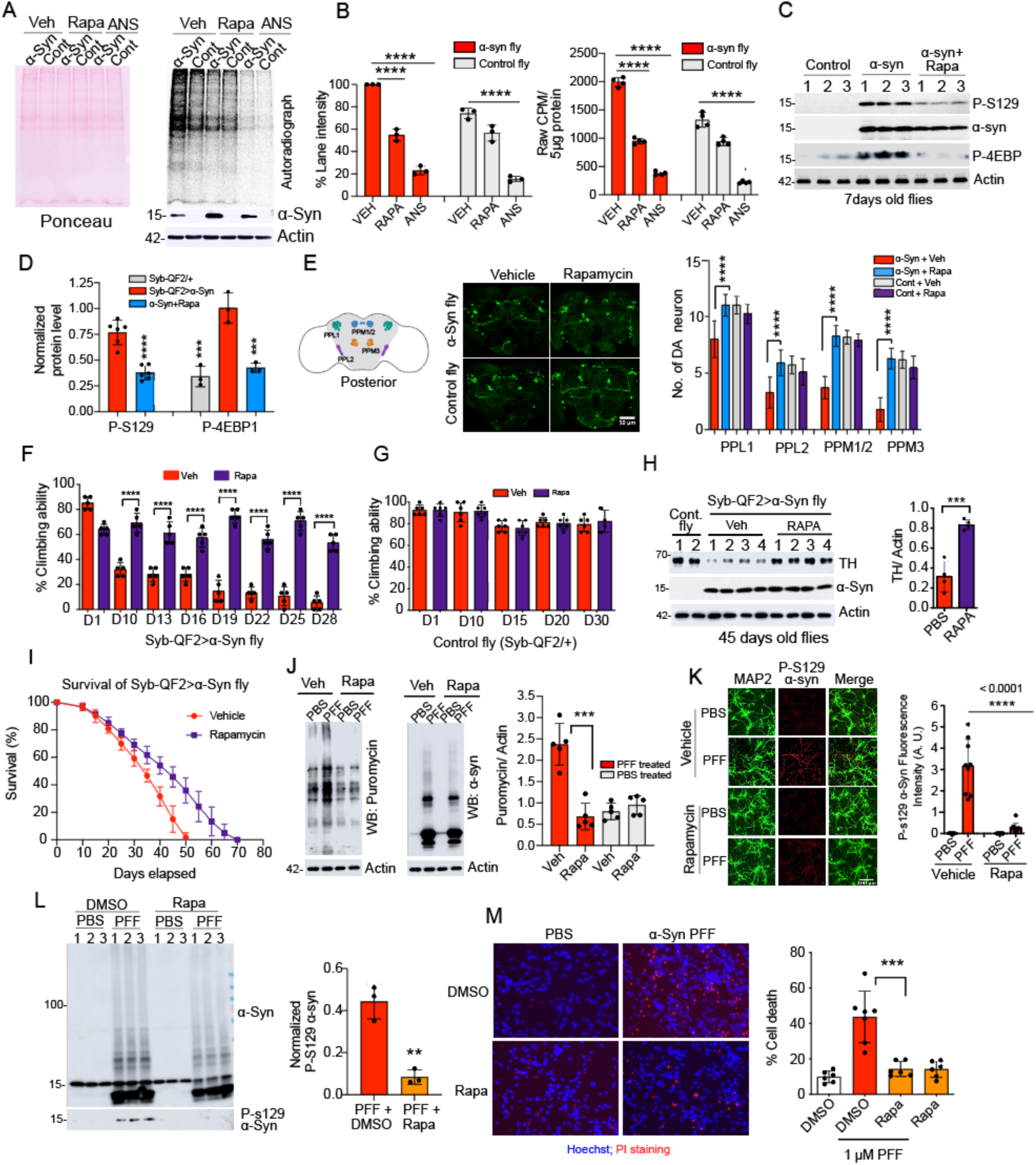
Pathogenic α-syn mediated increase in protein translation depends on mTORC1 kinase activity. (**A**) Supplementation of Rapamycin in regular fly food diminishes translation upregulation in α-syn flies (TH Gal4>α-syn). Autoradiograph shows S35 incorporation in fly brains (n=3). (**B**) The bar graphs show quantification of S35 intensity in the autoradiograph and quantification of scintillation counts of 5 μg total protein in each group (n=3). (**C**) Immunoblots show rapamycin treatment reduces α-syn pathology (P-S129 α-syn phosphorylation) and 4E-BP phosphorylation in Syb-QF2>α-syn flies. (**D**) The graph shows quantification of P-S129 and 4E-BP phosphorylation in Figure 3C (n=3). (**E**) Representative confocal images show whole brain TH immunostaining of posterior DA neuron clusters in control (Syb-QF2/+) and Syb-QF2>α-syn flies. Scale = 50 μM. The bar graph shows quantification of TH positive neurons in the indicated posterior DA neuron clusters (n=8). (**F**) The bar graph shows percent climbing ability of vehicle and rapamycin treated Syb-QF2>α-syn flies. The climbing abilities were assessed up to 30 days (n=5). (**G**) Rapamycin has no adverse effect on motor function in control flies. The graph shows percent climbing ability of rapamycin treated control flies (n=6). (**H**) Immunoblots show TH, α-syn and actin protein levels in the brain tissue of rapamycin treated flies. The bar graph shows quantification of normalized TH level (n=3). (**I**) Survival curve shows increased life span of rapamycin treated Syb-QF2>α-syn flies (n=4). (**J**) Rapamycin treatment (0.5 μM) diminishes α-syn PFF induced translation in primary cortical neurons. The immunoblot was probed for puromycin, actin and α-syn protein. The graph shows quantification of normalized new protein synthesis with actin level, n=4. (**K**) Rapamycin treatment prevents α-syn PFF induced pathology in primary cortical neurons. Confocal images show P-S129 α-syn immunostaining in cortical neuron. Graph shows the quantification of P-S129 α-syn signal intensity (n=9). Scale = 100 μM. **(L)** The immunoblots show protein level of α-syn and p-S129 α-syn in α-syn PFF and rapamycin treated primary neurons. The bar graph shows quantification of normalized P-S129 α-syn levels (n=3). (**M**) Representative images of Hoechst and PI staining show rapamycin rescues neurotoxicity in α-syn PFF treated cortical neurons. The bar graph shows percent cell death in α-syn PFF and rapamycin treated neurons (n=3). For figures 3B, 3F, 3H, 3J and 3L statistical significance was measured by unpaired two-tailed t test. Data are expressed as mean ± SEM; ns, not significant and *p ≤ 0.05, **p ≤ 0.01, and ***p ≤ 0.001. For the figures 3A, 3D, 3E and 3K statistical significance was measured by One-way ANOVA, data are expressed as mean ± SEM, *p < 0.05, **p < 0.01, ****p < 0.0001.

The *Drosophila* Syb-QF2> α-syn model, which exhibits robust pathologic α-syn induced degeneration (*16*), showed a significant increase in P-S129 α-syn and d4EBP phosphorylation that was reduced by feeding the flies, the mTOR inhibitor, rapamycin (Figures 3C and 3D). Dopamine neuron loss and locomotion defects in the Syb-QF2> α-syn flies were also reduced and lifespan was increased by feeding rapamycin (Figures 3E, 3F, 3H, 3I and Video S1). No adverse effect of feeding rapamycin was observed in control Syb-QF2/+ flies (Figure 3G and Video S2). In α-syn PFF treated primary cortical neurons, rapamycin showed a significant reduction in the α-syn PFF increased protein translation (Figure 3J). Rapamycin also significantly reduced α-syn PFF induced P-S129 α-syn levels (Figure 3K and 3L) and it reduced α-syn PFF neurotoxicity (Figure 3M). These findings suggest that pathologic α-syn mediated increase in translation depends on mTOR and that rapamycin neutralizes the increased protein translation and rescues PD pathology in *Drosophila* and mouse cortical neurons.

### The mTOR Inhibitor, Rapamycin is Neuroprotective in the Intrastriatal α-syn PFF Mouse Model

Next, we examined the effect of rapamycin in the intrastriatal α-syn PFF mouse model (*17*). Intrastriatal α-syn PFF mice were injected intraperitonially with rapamycin (6 mg/Kg; #R-5000, LC laboratories) or vehicle (10% PEG400, 10% Tween 80 in water) on the day of the intrastriatal α-syn PFF injection and every two days thereafter following the intrastriatal α-syn PFF injection for 30 days. Administration of rapamycin reduced P-S129 α-syn immunoreactivity throughout the brain of α-syn PFF injected mice (Figures S3A and S3B). Rapamycin also reduced S6K and 4E-BP1 phosphorylation throughout the brain that was induced by the intrastriatal α-syn PFF injection (Figures S3C-S3F). Another group of mice received an intrastriatal α-syn PFF injection and were fed rapamycin the following day for 6 months. Rapamycin significantly reduced the loss of both TH and Nissl positive neurons as determined using unbiased stereology (Figure 4A). Rapamycin feeding also led to a significant reduction in P-S129 α-syn (Figures 4B and 4C) and it reduced S6K and 4E-BP1 phosphorylation throughout the brain at six months in the α-syn PFF injected mice (Figures 4D and 4E). Rapamycin had no effect on body weight in intrastriatal α-syn PFF and control injected mice (Figure 4F). Rapamycin rescued the behavioral deficits induced by intrastriatal α-syn PFF injection as assessed via the pole test (*18*) (Figures 4G and 4H) and by assessing forelimb and hindlimb grip strength (*18*) (Figures 4I and 4J). Immunoblot analysis of α-syn PFF treated primary cortical neurons confirms that rapamycin reduced S6K and 4E-BP1 phosphorylation (Figure 4K). Taken together these data support that the mTOR inhibitor rapamycin is neuroprotective in the α-syn PFF mice model.

**Fig. 4:**
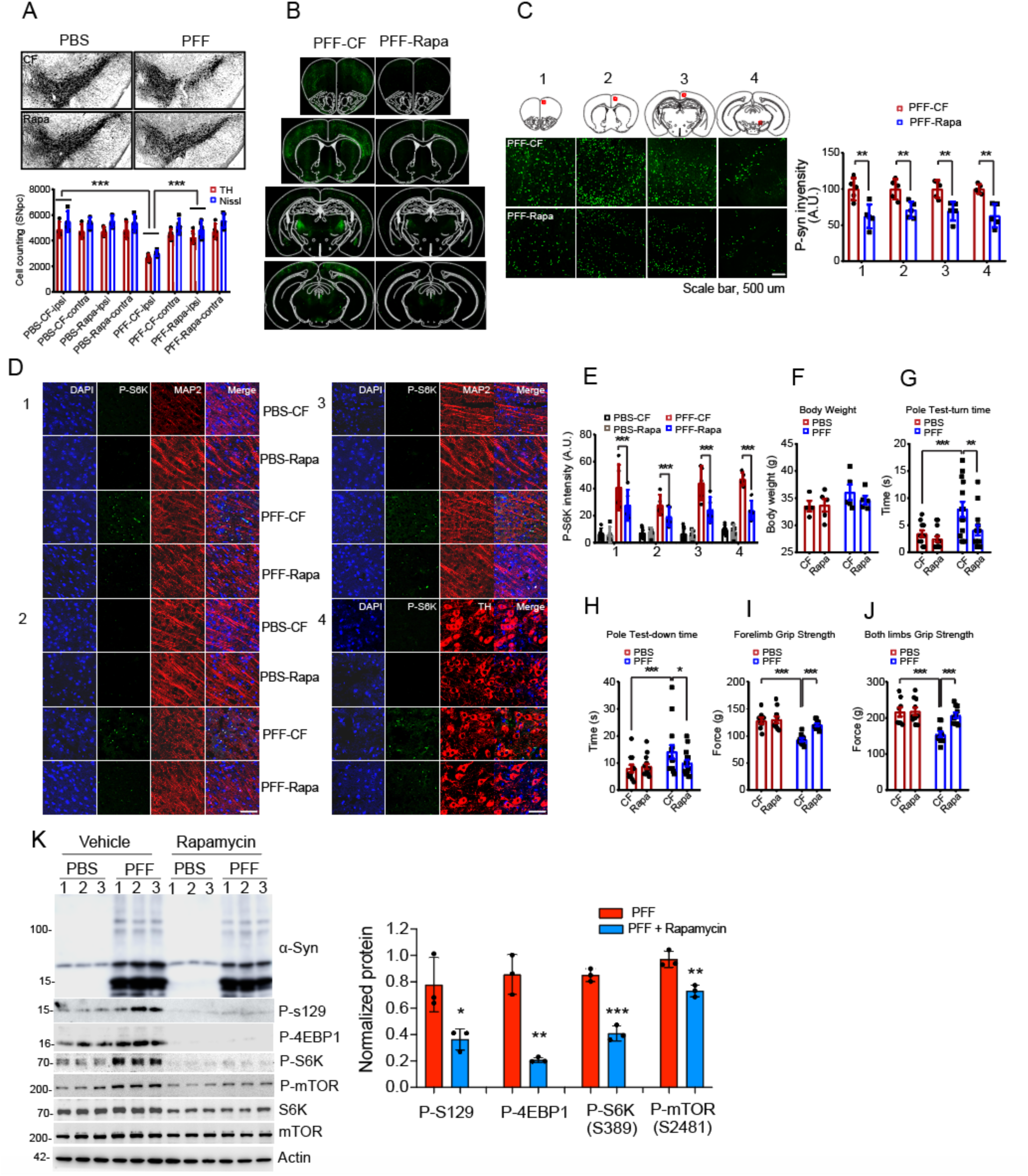
Rapamycin is neuroprotective in the α-syn PFF mouse model. (**A**) Representative images of TH immunostaining of the substantia nigra (SN) 6 months post intrastriatal α-syn PFF injection treated with rapamycin (Rapa) or control food (CF). The bar graph shows quantification of TH and Nissl positive cells in the SN (n=5). (**B**) Representative images P-S129 α-syn in whole brain coronal sections of different brain regions (n=5). (**C**) α-syn pathology in the zoom-in area of different brain regions. The bar graph shows quantification of P-S129 α-syn intensity in the zoom-in area (n=5). (**D**) Representative images of P-S6K (P-T389) immunostaining in the indicated brain regions. (**E**) The bar graph shows quantification of P-S6K signal intensity in Figure 4D (n=5). (**F**) The bar graph shows body weight of mice with CF and Rapa diet (n=5). **(G)** Pole test turn-time (n=5). **(H)** Pole test down times (n=5). (**I & J**) The bar diagrams show fore limb and both limbs grip strength respectively (n=5). (**K**) Rapamycin suppresses α-syn pathology and mTORC1 pathway in primary cortical neurons. Immunoblots show the levels of P-S129 α-syn, P-4EBP1, P-S6K, and P-mTOR (P-S2481) in treated and untreated neurons. The bar graph shows quantification of P-S129 α-syn, P-4EBP1, P-S6K, and P-mTOR (P-S2481) in the immunoblots (n=3). For the figures 4C and 4K statistical significance was measured by unpaired two-tailed t-test. Data are expressed as mean ± SEM; and *p ≤ 0.05, **p ≤ 0.01, and ***p ≤ 0.001. For the figures 4A and 4E-4J statistical significance was measured by One-way ANOVA, data are expressed as mean ± SEM, *p < 0.05, **p < 0.01, ****p < 0.0001.

### Increase Protein Translation Contributes to Neurotoxicity Induced by Pathologic α-syn

Consistent with mTOR activation, S6K is phosphorylated and activated in transgenic *Drosophila* models (Figures S4A and S4B). Since S6K phosphorylation by mTOR is linked to translation activation (*19*), we explored the possibility of whether specific inhibition of S6K could rescue PD pathology and neurodegeneration in the pathologic α-syn model (Syb-QF2> α-syn) (*16*). Indeed, feeding of the S6K inhibitor (PF-4708671) by adding to fly food significantly prevents the TH loss (Figure S4C) and motor deficits in the Syb-QF2> α-syn flies (Figure S4D and Video S5), while the S6K inhibitor had no substantial effect on control Syb-QF2/+ flies (Figure S4E and Video S6). Inhibition of S6K significantly prevented DA neuron loss and increased the lifespan of Syb-QF2>α-syn flies (Figure S4F and S4G). Inhibition of S6K also showed reduced α-syn PFF induced P-S129 α-syn immunoreactivity and toxicity in primary cortical neurons (Figures S4I and S4J). The syb-QF2>α-syn *Drosophila* model exhibited a significant increase in P-S129 α-syn and d4EBP phosphorylation (Figure 5A). Feeding syb-QF2>α-syn flies, the global protein synthesis inhibitor, anisomycin, significantly reduced enhanced P-S129 α-syn phosphorylation (Figure 5B) and it reduced TH loss as determined by immunoblot analysis (Figure 5C). Anisomycin also significantly reduced DA neuron loss and locomotion defects (Figures 5D, 5E, 5F and Video S3). Anisomycin feeding had no substantial effect in control Syb-QF2/+ flies (Figure 5G and Video S4). The lifespan of the Syb-QF2/+ *Drosophila* model was significantly increased by anisomycin feeding (Figure 5H). Anisomycin also reduced α-syn PFF induced P-S129 α-syn immunoreactivity (Figure 5I and 5J) and toxicity (Figure 5K) in primary cortical neurons. Taken together, these data suggest that increased protein translation contributes to the neurotoxicity induced by pathologic α-syn.

**Fig. 5:**
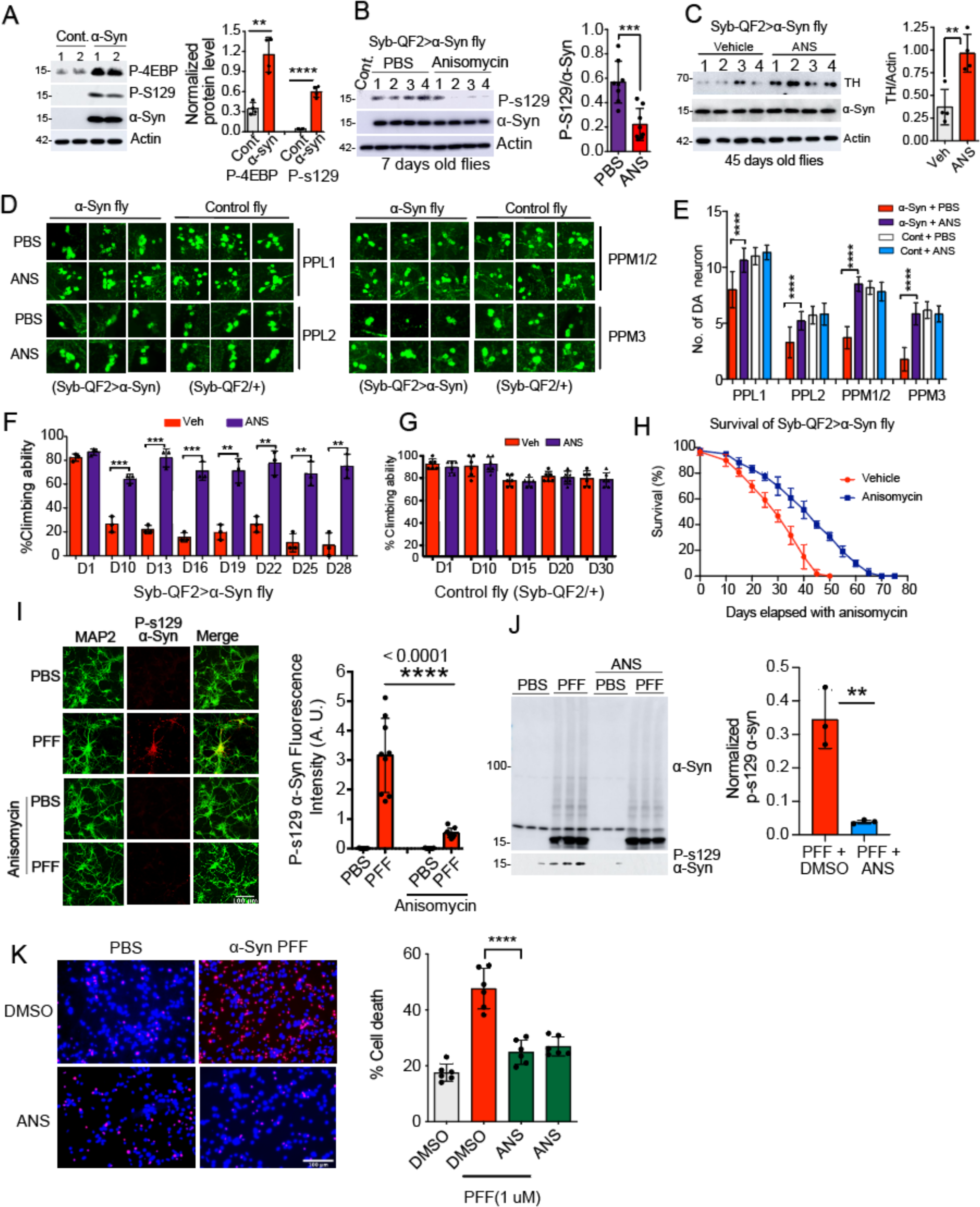
Anisomycin rescues α-syn pathology and neurotoxicity. (**A**) Immunoblots show significant α-syn phosphorylation (P-S129 α-syn) and phosphorylation of endogenous d4EBP in Syb-QF2>α-syn flies. The graph shows quantification of P-d4EBP and P-S129 α-syn levels, n=4. (**B**) Immunoblots shows P-s129 α-syn and bar graph shows quantification of normalized P-S129 α-syn level, n=8. (**C**) The immunoblots level of TH, α-syn and actin. The graph shows quantification of normalized TH level (n=4). (**D**) Representative confocal images show individual DA neuron clusters in control and α-syn transgenic flies. (**E**) Quantification of DA neurons in the indicated posterior DA neuron clusters (n=8). (**F**) The bar graph shows percent climbing ability of anisomycin treated α-syn transgenic fly flies (Syb-QF2>α-syn) (n=3). (**G**) The bar graph shows percent climbing ability of anisomycin treated control flies (Syb-QF2/+) (n=6). (**H**) Survival curve shows increased life span in anisomycin treated Syb-QF2>α-syn flies (n=4). (**I**) Representative confocal images of P-S129 α-syn immunostaining of PFF treated primary cortical neuron. The graph shows quantification of signal intensity of P-S129 α-syn (n=3). Scale = 100 μM. (**J**) Immunoblot shows α-syn pathology (P-S129 α-syn) in primary cortical neurons. The bar graph shows quantification of normalized P-S129 α-syn level (n=3). (**K**) Hoechst and PI staining shows anisomycin rescues neurotoxicity in α-syn PFF treated neurons. The bar graph shows quantification of cell death, n=6. For the figures 5A, 5B, 5C and 5F statistical significance was measured by unpaired two-tailed t test. Data are expressed as mean ± SEM; ns, not significant and *p ≤ 0.05, **p ≤ 0.01, and ***p ≤ 0.001. For the figures 5E, 5I, 5J and 5K statistical significance was measured by One-way ANOVA, data are expressed as mean ± SEM, *p < 0.05, **p < 0.01, ****p < 0.0001.

### Pathologic α-syn Binds TSC2 and Disassembles the TSC1-TSC2 Complex

Since pathologic α-syn interacts with the TSC2 protein (see Figure 1E and 1F) and the TSC1-TSC2 complex plays an important role in mTORC1 activity, the consequences of this interaction on TSC1-TSC2 complex stability using Single Molecule Pulldown (SiMPull) assays were evaluated. The SiMPull assay utilizes a combination of immunoprecipitation and visualization of single molecules by fluorescence imaging to study interaction dynamics within protein complexes (*20-22*). SiMPull was used to pulldown the TSC1-TSC2 complex on glass slides from cell extracts by capturing either FLAG tagged TSC2 or Myc tagged TSC1. TSC1 and TSC2 were captured by biotinylated anti-Myc and ant-FLAG on neutravidin coated glass surfaces and visualization was accomplished using Alexa-647 tagged anti-rabbit secondary antibodies (Figure 6A). The addition of pathologic α-syn PFF resulted in a dose dependent disassembly of the TSC1-TSC2 complex, and quantification revealed that 0.5 nM α-syn PFF resulted in the dissociation of 50% of the TSC1-TSC2 protein complex (Figure 6A). We examined the effect of monomeric α-syn protein in the same experimental set up and observed no significant change in TSC1-TSC2 complex stability (Figure S5A). Next, we asked whether pathologic α-syn has any adverse effect on mTORC1 complex stability. Our data revealed that mTORC1 complex is stable, even at high concentration of α-syn PFF (5000 nM) (Figure 6B). In a similar manner, monomeric α-syn shows no effect on mTORC1 complex stability (Figure S5B). Aβ oligomers and Tau PFF do not disassemble the TSC1-TSC2 complex (Figure 6C) suggesting the destabilization of the TSC1-TSC2 protein complex is specific for pathologic α-syn. We examined the effect of α-syn PFF on PKA complex previously used as a control in the SiMPull assay (*20-22*) and found no effect on PKA complex (Figure S5C). We further validated the effect of α-syn PFF in the disassembly of TSC1-TSC2 complex using conventional immunoprecipitation (IP) and western blot. Consistent with the SiMPull data, the TSC complex is disassembled by α-syn PFF in a dose dependent manner (Figure 6D).

**Fig. 6:**
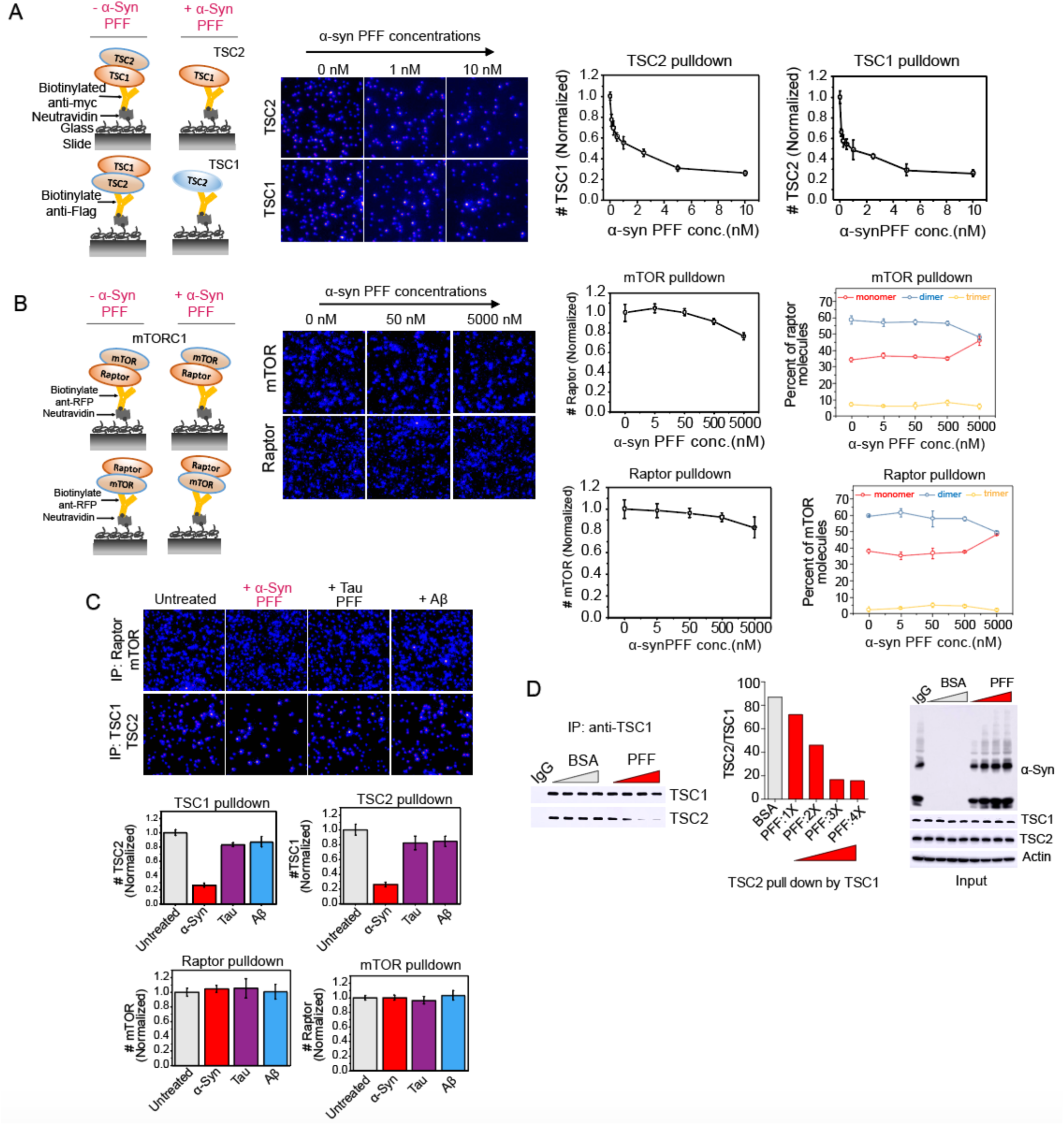
Pathogenic α-syn destabilizes the TSC1-TSC2 complex. (**A**) Schematic shows TSC1-TSC2 SiMPull assay design in absence and presence of pathogenic α-syn. Representative images show single molecule pulldown of TSC1 and TSC2 in the presence of different concentrations of α-syn PFF. The line graphs show normalized level of TSC1 and TSC2 molecules pulled down by TSC2 and TSC1 respectively (n=3). (**B**) Schematic shows SiMPull assay design for the mTORC1 complex in absence and presence of pathogenic α-syn. Representative images show single molecule pulldown of mTOR and Raptor in presence of different concentrations of recombinant α-syn PFF. The middle line graphs show normalized level of mTOR, and raptor molecules pulled down by raptor and mTOR respectively. The right line graphs show different species of raptor and mTOR molecules in presence of α-syn PFF (n=3). (**C**) Representative images show single molecules of mTOR and TSC2 pulled down by raptor and TSC1 respectively in the presence of α-syn PFFs, Tau PFFs and Aβ oligomers. The bar graphs show normalized levels of mTOR, raptor, TSC1 and TSC2 single molecules in SiMPull assay in presence of toxic aggregates (n=3). (**D**) Conventional immunoprecipitation and western blots show pathologic α-syn disrupts the TSC1-TSC2 complex. IP immunoblots shows pulldown of TSC2 by TSC1 in presence of increasing concentrations of α-syn PFF. Actin was used as a loading control. The bar graph shows quantification of normalized TSC2 protein levels in the IP. In the IP 0.25 μM PFF concentration was used as 1X.

### Constitutive Overexpression of 4E-BP1 and TSC2 Rescues Motor Deficits and PD Pathology in the **α-syn Fly Model**

Activated mTOR phosphorylates 4E-BP1, which leads to inhibition of 4E-BP1’s repression of 5’cap-dependent translation resulting in the initiation of 5’cap-dependent translation (*23, 24*). Since pathologic α-syn activation leads to 4E-BP1 phosphorylation via mTOR activation, the effect of constitutive overexpression of active 4E-BP1 was evaluated. Constitutive overexpression of 4E-BP1 in Syb-QF2>α-syn *Drosophila* model rescued the severe motor deficit of the Syb-QF2> α-syn flies (Figure S6A and Video S8). Double transgenic flies showed significantly less P-S129 α-syn levels (Figure S6B). Both TH and DA neuron loss were also rescued by 4E-BP1 overexpression (Figure S6C-S6E). Mitochondrial degeneration has been implicated in α-synucleinopathy (*13, 25*) and α-syn transgenic flies show abnormal mitochondrial morphology (*16*). Since mTORC1 and 4E-BP1 play important role in mitochondrial health (*26, 27*), mitochondrial morphology using Transmission Electron Microscopy (TEM) was examined. In Syb-QF2> α-syn flies, mitochondria are enlarged which was essentially normalized in double transgenic flies comparable to Syb-QF2/+ control flies (Figures S6F-S6I). In the primary α-syn PFF cortical neuronal model, 4E-BP1 overexpression by Lentivirus significantly reduces the enhanced protein translation (Figure S6J) and significantly reduced P-S129 α-syn levels and neurotoxicity (Figures S6K and S6L).

TSC2 constitutively downregulates mTORC1 activity by GTP hydrolysis of the Rheb GTPase (*28*). Since pathologic α-syn directly binds TSC2 and disassembles the TSC1 and TSC2 complex, we hypothesized that constitutive overexpression of TSC2 will result in 1) neutralization of the effect of pathologic α-syn leading to 2) the suppression of aberrant mTORC1 signaling. TSC2 overexpression in the Syb-QF2>α-syn flies significantly rescues the severe motor deficit in Syb-QF2>α-syn flies (Figure 7A and Video S7). TSC2 overexpression also significantly reduced P-S129 α-syn levels (Figure 7B) and TH loss as determined via immunoblot analysis (Figure 7C). Pathologic α-syn interacts with TSC2 in Syb-QF2>α-syn flies (Figure 7D). Confocal z-stack projection imaging for quantification of dopamine neurons in double transgenic Syb-QF2>α-syn:TSC2 fly brains showed significant neuroprotection in the posterior medial clusters (Figure 7E). Mitochondrial morphology and abundance in Syb-QF2>α-syn:TSC2 fly brains are comparable to control flies (Figure 7F-7I). The increase in protein translation and P-S129 α-syn levels were rescued by overexpression of TSC2 in α-syn PFF treated primary cortical neurons (Figures 7J and 7K). TSC1 is an HSP-90 co-chaperone and protects TSC2 from HERC1-mediated ubiquitination and degradation (*29*). Since pathologic α-syn destabilizes the TSC1-TSC2 complex, the levels of TSC1 and TSC2 were assessed in α-syn PFF treated primary neurons and postmortem PD brains. α-Syn PFF treatment caused a significant decrease in TSC1 and TSC2 protein levels (Figure 7L). In human substantia nigra (Figure 7M) and the Lewy body rich cingulate and inferior parietal cortex (Figure 7N) there was a significant reduction in TSC1-TSC2 protein levels in the setting of increased P-S129 α-syn levels (Figures 7M and 7N). There were no significant change TSC1-TSC2 protein levels and P-S129 α-syn was not detectable in the cerebellum, a relatively unaffected brain area in PD (Figure 7O).

**Fig. 7:**
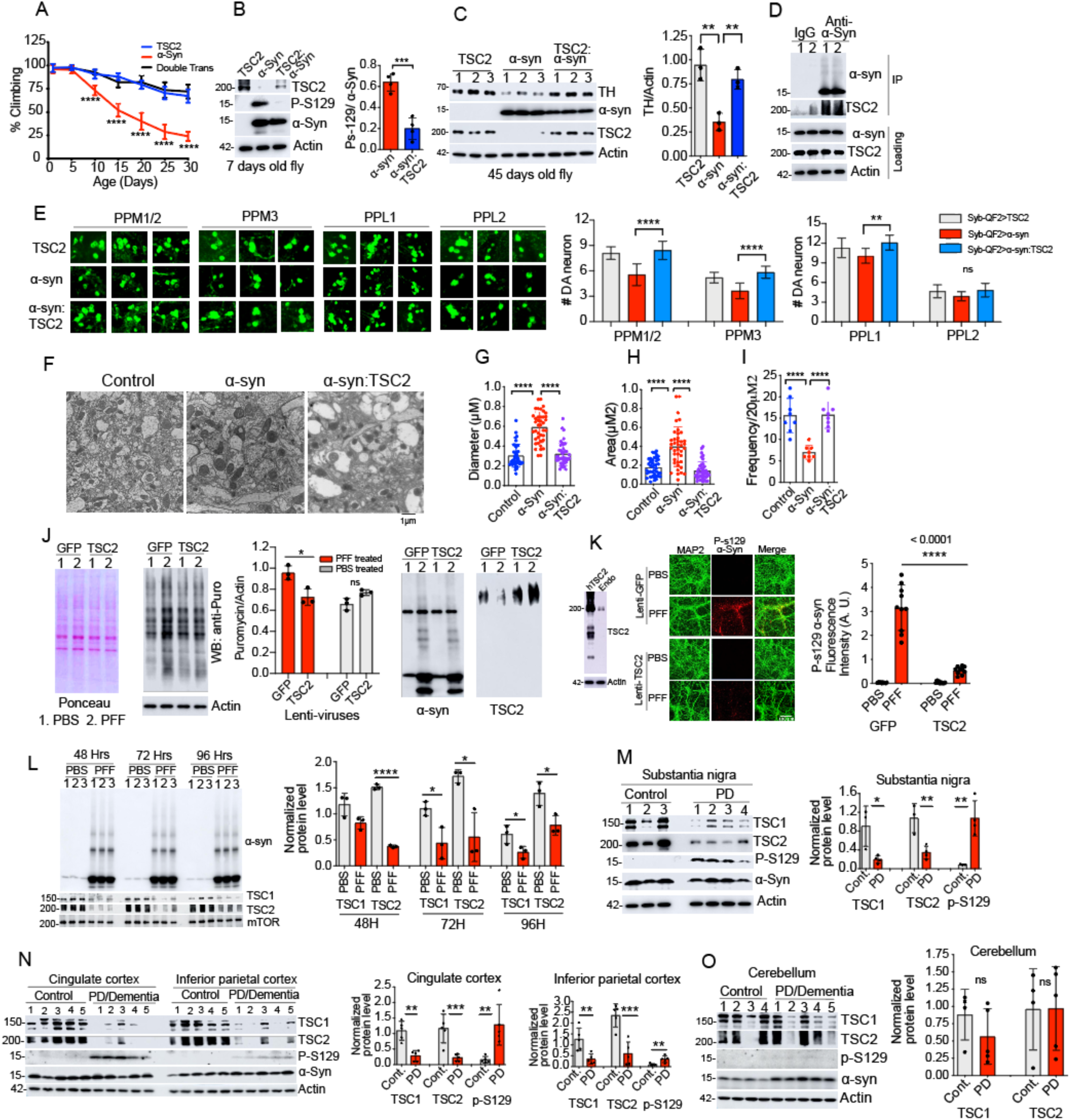
Constitutive overexpression of 4E-BP1 and TSC2 rescues motor deficits and PD pathology in the α-Syn fly model. (**A**) Line graph shows quantification of percent climbing abilities of young and old transgenic flies. Climbing abilities were monitored for 30 days (n=5). (**B**) Immunoblots show level of P-S129 α-syn in transgenic flies. The bar graph shows quantification of P-S129 level in the immunoblot (n=4). (**C**) Immunoblots show protein levels of TH, α-syn, TSC2 and actin in fly brain homogenates. The graph shows the quantification of normalized TH level (n=3). (**D**) Co-Immunoprecipitation shows pulldown of α-syn and TSC2 from fly brain homogenates. (**E**) Representative confocal images show individual DA neuron clusters in 45 days old transgenic flies. The bar graphs show quantification of DA neuron counts in the indicated clusters (n=7). (**F**) Transmission Electron Microscopic (TEM) images show mitochondrial size and frequency in transgenic flies. Scale = 1 μM. (**G-I**) Quantification of mitochondrial diameter, area, frequency in the control (Syb-QF2/+) and transgenic flies. (**J**) The blots from SUnSET assay show ponceau and probing for puromycin, α-syn and TSC2. The bar graph shows quantification of new protein synthesis (n=3). (**K**) The immunoblot shows expression of TSC2, representative confocal images show P-S129 α-syn and MAP2 staining in α-syn PFF treated neurons. Scale = 100 μM. The graph shows quantification of P-S129 α-syn intensity. (**L**) Immunoblots of TSC1, TSC2 and mTOR protein levels in α-syn PFF treated primary cortical neurons at different time points. The graph shows quantification of TSC1, TSC2 and mTOR protein levels (n=3). (**M**) The immunoblots show protein levels of TSC1, TSC2, P-S129 α-syn, total α-syn and actin in substantia nigra of post-mortem PD brains. The graph shows quantification of TSC1, TSC2 and P-S129 α-syn protein level (n=3). (**N**) The immunoblots show protein levels of TSC1, TSC2, P-S129 α-syn, total α-syn and actin in cingulate and inferior parietal cortex from post-mortem PD and control brains. The bar graph shows quantification of normalized TSC1, TSC2 and P-S129 α-syn protein level (n=5). (**O**) The Immunoblots show protein levels in cerebellum and the graph shows quantification of TSC1 and TSC2 protein levels (n=3). For the figures 7J, 7L-7O statistical significance was measured by unpaired two-tailed t test. Data are expressed as mean ± SEM; ns, not significant and *p ≤ 0.05, **p ≤ 0.01, and ***p ≤ 0.001. For the figures 7A-C,7F, 7G-I and 7K statistical significance was measured by One-way ANOVA, data are expressed as mean ± SEM, *p < 0.05, **p < 0.01, ****p < 0.0001.

## DISCUSSION

The major finding of this study is that pathologic α-syn binds TSC2 and destabilizes the TSC1-TSC2 complex leading mTORC1 pathway activation and enhanced protein translation. In support of these observations, we found aberrant mTORC1 pathway activation in fly and mouse pathologic α-syn models of neurodegeneration and in human postmortem PD brains. Our translation data that included a cell free IVT assay, a neuron based SUnSET assay and in vivo S35 pulse labelling, supported mTOR dependent and enhanced mRNA translation. Both genetic and pharmacological inhibition of translation and mTORC1 activity rescued neurotoxicity in the intrastriatal α-syn PFF mouse model, α-syn PFF treated primary cortical neurons and *Drosophila* α-syn transgenic modes. Together, our data uncover a molecular mechanism by which pathologic α-syn directly binds to TSC2 and destabilizes the TSC1-TSC2 complex and triggers aberrant mTORC1 activation, which results in enhanced mRNA translation and subsequent neurodegeneration.

Another study that used APEX labelling in neurons and mass-spectrometry also identified connections of α-syn with mRNA metabolism and translation where α-syn was reported to directly associate with initiation-elongation factors and mRNA binding proteins (*30*). This earlier study, which focused on monomeric α-syn, reported that α-syn suppressed mRNA translation. Consistent with the suppression of mRNA translation, we report that monomeric non-pathologic α-syn suppresses protein translation. This contrast with our findings that pathologic α-syn enhances protein translation. These findings taken together suggest that monomeric non-pathologic suppresses protein translation while pathologic α-syn enhances protein translation Our in vitro and in vivo functional and biochemical data suggest that destabilization of the TSC1-TSC2 complex by pathologic α-syn leads to several important molecular events. First, is the activation of mTOR kinase. Second, is the phosphorylation and inactivation of 5’cap-dependent translation repressor 4E-BP1. Third, is mTOR kinase phosphorylation of S6K. Fourth, is enhanced mRNA translation. These events are critically linked to neurodegeneration mediated by pathologic α-syn since restoration of TSC2 levels, inhibition of mTOR, expression of 4E-BP1, inhibition of S6K or inhibition of protein translation exhibited neuroprotection against pathologic α-syn. Consistent with our findings, prior studies have shown that 4E-BP1 overexpression provides neuroprotection against misfolded protein stress and α-syn mediated neurodegeneration (*26*). Moreover, activation of 4E-BP1 by rapamycin treatment is sufficient to rescue dopamine neuron loss and motor deficits in *Drosophila* PINK1 and parkin models of DA neuron loss (*7*). Similar to our findings in PD, in Huntington’s disease, pharmacological inhibition of mTOR by rapamycin is neuroprotective and this effect is attributed to the activation of 4E-BP1 and inhibition of protein synthesis in Huntington’s disease (*31*).

In summary we establish a connection between pathologic α-syn and deregulation of mRNA translation in PD through activation of mTORC1 via destabilization of the TSC1-TSC2 complex. Strategies aimed at reducing mTORC1 activity in PD may have therapeutically utility by restoring the imbalance in mRNA translation.

## MATERIALS AND METHODS

### EXPERIMENTAL MODEL AND SUBJECT DETAILS

#### Animals

All animal procedures following the guidelines of Laboratory Animal Manual of the National Institute of Health Guide to the Care and Use of Animals and upon approval of the Johns Hopkins Medical Institute Animal Care and Use Committee. C57BL/6J male mice (Jackson Laboratory) were used for intrastriatal α-syn PFF injections. These α-syn PFF injected mice were divided into two groups: one group of mice given rapamycin intraperitonially for one month and another group of mice were given microencapsulated rapamycin (eRapa) supplemented in regular mouse diet (*32*). Human alpha-syn(A53T) transgenic G2-3 lines are available from Jackson Laboratory (*13*) were aged and late symptomatic mice were used for biochemical analysis and isolation of endogenous A53T α-syn aggregates. Genotyping of A53T mice was carried out according to the Jackson Laboratory Protocol 25347.

#### Drosophila

D. melanogaster QUAS-SNCA flies were a kind gift from Dr. Mel B. Feany (Harvard Medical School) (*16*). D. melanogaster Syb-QF2 flies were a kind gift from Dr. Christopher J Potter (Johns Hopkins Medicine). D. melanogaster QUAS-4EBP1 and D. melanogaster QUAS-TSC2 flies were generated in this study. D. melanogaster UAS-SNCA flies were obtained from Bloomington Drosophila Stock Center (BDSC) of Indiana University.

#### Cell culture

HEK293FT cell line was obtained from ThermoFisher Scientific and used for in situ labelling in A53T α-syn-APEX mass spectrometry experiment. This cell line was routinely maintained in Dulbecco’s Modified Eagle Medium (DMEM) with 10% (v/v) bovine serum (FBS) and 1% Penicillin-Streptomycin, at 37^°^C with a 5% CO2 atmosphere in a humidified incubator. Wild type mouse primary neuron cultures were prepared from embryonic day E15 pups, cultured, and maintained in neurobasal media with B27 supplement (Gibco) and 1% Penicillin-Streptomycin in a 37°C humidified incubator.

#### Recombinant α-syn PFFs

Recombinant human and mouse α-syn PFFs were prepared as previously described (*33*). Recombinant α-syn proteins were expressed in BL2-competent E. coli and bacteria were grown overnight in terrific broth at 37^°^C with continuous agitation. After overnight incubation bacterial pellets were collected and resuspended in ice cold high-salt buffer (1 mM EDTA, 1mM PMSF, and protease inhibitors) followed by sonication on ice and boiling at 100°C for 15 min. After centrifugation at 6000 x g, 4^°^C supernatants were collected and dialyzed ((10 mM Tris (pH 7.6), 50 mM NaCl, 1 mM EDTA) at room temperature for an hour and the dialysis cassettes were transferred to fresh dialysis buffer of overnight dialysis at 4^°^C. The dialyzed proteins were concentrated with centrifugation filter unit (3.5 kDa cutoff, Amicon). The protein samples were loaded into FPLC for column separation using Superdex 200 SEC column. The protein fractions were buffer exchanged (10 mM Tris, pH 7.6, 25 mM NaCl, 1 mM EDTA) and performed anion-exchange chromatography (Hi-Trap Q HP) for further purification. The final protein fractions were collected, and buffer exchanged (50 mM NaCl) to obtain purified α-syn monomeric proteins. The purified proteins were made endotoxin free using the Toxineraser endotoxin removal kit (Genscript, Piscataway, NJ). For fibrillation, monomeric α-syn proteins in PBS (5 mg/mL) were incubated at 37^°^C with continuous agitation (1000 rpm) for one week and fibrils are stored at -80^°^C. For intrastriatal injection, α-syn PFFs were thawed, diluted in PBS (2.5 mg/mL), and sonicated in ice at 30% amplitude for 120 seconds (1s on/off) using Branson Digital Sonifier (Danbury, CT, USA).

## METHODS DETAILS

### Reagents

Key resources and reagents are list in Table S3.

### In vitro translation (IVT) assay

Translation extract was prepared from human neuroblastoma (SHSY-5Y) or HEK293 cells grown in 175 cm^2^ flask to achieve ∼80% confluence. The growing cells were harvested using trypsinization and neutralization with complete DMEM media. The cell pellets were washed with 10 mL PBS to remove residual media from the cell pellet. After PBS wash, the cell pellets were kept in ice and lysed immediately in freshly made translation extract lysis buffer (30 mM HEPES-KOH, 100 mM potassium acetate, 2 mM magnesium acetate, 5 mM DTT, 1 mg/mL Pefabloc SC). The cell pellet from two 175 cm^2^ flasks was lysed in 500 μL lysis buffer. The lysis was performed using mechanical homogenizer and Corning PYREX Tissue Grinder in ice. Total twenty strokes were applied to homogenize the cell pellet. The lysates were centrifuged at 12500 rpm for 20 mins at 4°C. After centrifugation the clear cytoplasmic fractions were collected from the top of the centrifuge tube. The cytoplasmic fractions are aliquoted and stored at -80°C.

### Translation assay design

Translation reaction was assembled as described previously (*34*). A 25μL translation reaction consisting of 50 ng luciferase reporter, 40% (v/v) translation extract and 60% (v/v) translation mix (16 mM HEPES-KOH at pH 7.4, 100 μM complete amino acid mix, 50 mM potassium phosphate, 2.5 mM magnesium acetate, 100 μM spermidine, 250 μg/mL yeast tRNA, 80 μg/mL creatine kinase, 20 mM creatine phosphate, 800 μM ATP, 100 μM GTP) were incubated at 30° C thermo cycler for 30 minutes. Firefly luciferase activity was measured using Promega luciferase assay system.

### Drosophila S35 assay

S35 assay in *Drosophila* was performed as described previously (*5*). α-Syn transgenic and age matched control flies (5 days old) were fed with radioactive S35 (Perkin Elmer) for 10 hours. 10 μCi S35 in 1 mL PBS (supplemented with 2% sucrose) applied to wet Kim wipes in 9 cm empty Drosophila vials (Genesee Scientific, cat no. 32–113). In each fly vial 20 flies were housed for S35 feeding. After S35 feeding flies were snap frozen in liquid nitrogen and heads were separated from body by vortexing and sieving. Equal number of fly heads were lysed in 1X RIPA buffer (supplemented with protease inhibitor) using mechanical homogenizer. Fly brain homogenates were collected in clean Eppendorf tube and protein concentrations were measured using BCA (Thermo) kits. For autoradiograph, total 10 μg protein was loaded per sample in 4-20% Tris-Glycine grading PAGE (Invitrogen), transferred to PVDF membrane (BIO-RAD) overnight, stained for Ponceau (Ponceau S solution, SIGMA), then the PVDF membrane was air dried and exposed to hot plate and autoradiographed in Typhoon FLA 9500 (GE Healthcare) phospho imager. For direct scintillation counts total of 5 μg protein per sample was added to scintillation fluid and counts were measured in LS 6500 Scintillation Counter (Beckman Coulter).

### Direct autoradiography of fly heads

Transgenic and age matched control flies (5 days old) were fed with radioactive s35 (Perkin Elmer) for 10 hours. 10 μCi s35 in 1 mL PBS (supplemented with 2% sucrose) applied to wet Kim wipes in 9 cm fly vial. Total 20 flies were housed in each fly vials for s35 feeding. After feeding flies were snap frozen in liquid nitrogen and heads were separated from body by vortexing and sieving. The fly heads were placed on clean white paper, three heads were placed in each squire and the heads were fixed on the paper surface using adhesive plastic wrapper to prevent moving. The fly heads were autoradiographed in Typhoon FLA 9500 (GE Healthcare) phospho imager.

### SUnSET assay

The SUnSET assay was performed as previously described with minor changes (*12*). Primary cortical neurons were prepared from E15 pups and cortices from each pup were lysed and plated using neural basal medium. Mature cortical neurons (∼7 days old) were treated with human α-synuclein Pre-Formed Fibril (α-Syn PFF) in 5 μg/mL concentration in 12 wells culture plate for 15 hours. After PFF treatment the neurons were treated with Puromycin (Gibco) in 10 μg/mL concentration for 20 mins. After puromycin pulse the neurons were treated with cycloheximide (100 μg/mL) to block new protein synthesis. For western blot the PBS washed harvested cells were lysed in RIPA buffer and homogenized using mechanical homogenizer using 20 strokes. The clears lysates were collocated in Eppendorf and protein concentrations were estimated to run equal amount of protein for each sample. For each sample total 10 μg protein was loaded for western blot. To quantify new protein synthesis, the western blots were developed using anti-puromycin mouse Ab (Millipore) and quantifications were normalized by respective actin level. Puromycin immunohistochemistry was performed to visualize the incorporation of puromycin in the newly synthesized proteins. For puromycin histochemistry, primary neurons were grown and cultured on glass coverslip in 24 wells plate, α-Syn PFF treatment and puromycin pulse were given as above mentioned for puromycin western blot experiment. The cycloheximide treated neurons were washed with PBS and fixed with 300 μL cold 4% PFA by incubating at room temperature for 15 mins. The neurons on the coverslip were washed with 500 μL PBS for three times (5 mins each). After PBS wash primary antibodies were added in 1xPBS with 5% normal goat serum and 0.3% Triton-X in 250 μL volume for each coverslip. The primary antibodies were added in 1:1000 dilution for anti-puromycin (Mouse) and in 1:500 dilution for anti-NeuN (Rabbit) antibody and incubated overnight at 4°C shaker incubator. After primary antibody incubation the coverslips were washed with 1xPBS plus 0.3% Triton-X for 30 mins. The secondary antibodies (Alexa 488-anti Rb/Ms NeuN (1:1000) + Alexa 555-anti Ms/Rb p-a-syn (1:1000) were added and incubated at 37°C for 60 mins. After secondary antibody incubation the coverslips were washed for 30 mins and mounted on glass slides for confocal microscopy.

### Biotinylated α-Syn-PFF pulldown from mouse striatum

Three months old C57BL6 male mice were used for striatal injections of biotinylated human α-synuclein PFF. Mice were injected with 10 μg α-Syn PFF into both hemispheres. To control this experiment, C57BL6 male mice were injected with non-biotinylated human α-Syn PFF. After 5 days of injection, both mice were sacrificed and striatal tissues were isolated and lysed in 20 mM HEPES buffer (20 mM HEPES pH 7.4, 150 mM NaCl, 2.5 mM MgCl_2_, 0.1% Triton X-100, 0.1 % NP-40). Striatal tissues were homogenized in 800 μL lysis buffer using mechanical homogenizer and Corning PYREX Tissue Grinder in ice. Total twenty strokes were applied to homogenize the brain tissues. The homogenates were centrifuged at 12,000 rpm for 15 mins at 4°C and the clear lysates were collected in clean Eppendorf tubes. For streptavidin immunoprecipitation, 200 μL streptavidin M-280 Dynabeads (ThermoFisher) were washed and incubated with brain lysates for 4 hours at 4° C with continuous rotation. The immunoprecipitated beads were washed 5 times. Two sets of immunoprecipitations were performed. One pair of mice was used for mass spectrometry analysis and another pair was used for western blot analysis.

### In situ labelling of α-Syn binding partners using A53T α-Syn-APEX2 assay in HEK293 cell

In situ biotin labelling assay was performed as described by Lam et al. (*35*) with minor changes. In A53T-α-Syn-APEX2 construct, APEX2 was tagged to the C-terminal of A53T α-Syn under the CMV promoter. HEK293 cells were grown in complete DMEM and A53T α-Syn-APEX2 plasmid was transfected using Xtreme transfection reagent (Thermo). After 24 hours of HEK293 transfection, 500 mM biotin tyramide dissolved in Optimem was added to the transfected cells and incubated for 30 min at 37°C. To activate APEX2 enzyme activity, H_2_O_2_ was added at 1 mM final concentration for exactly 1 min at room temperature. Following H_2_O_2_ treatment the cells were washed twice with PBS containing quenching reagents (5 mM Trolox, 10 mM ascorbic acid and 10 mM NaN3) for 30s followed by another two washes with PBS. To control the immunoprecipitation, A53T α-Syn-APEX2 transfected cells were used without H_2_O_2_ treatment. The cells were homogenized in 20 mM HEPES buffer (20 mM HEPES pH 7.4, 150 mM NaCl, 2.5 mM MgCl_2_, 0.1% Triton X-100, 0.1% NP-40) and biotinylated protein complexes were Immunoprecipitated using streptavidin M-280 Dynabeads (ThermoFisher). Two sets of immunoprecipitations were performed. One set was used for mass spectrometry and another set was used for western blot analysis.

### Sample preparation for Mass Spectrometry

Captured biotinylated proteins using APEC protocol from HEK293T cells (*36*) and Immunoprecipitated samples from mouse brain tissue samples were boiled in 4X LDS sample buffer (Life Technologies) supplemented with 100 mM dithiothreitol or 200 mM beta-mercaptoethanol. These samples were loaded on Novex 4 to 20% Tris-Glycine protein gel (Thermo Fisher Scientific). Following the successful electrophoretic steps, gels were stained using a dye, Coomassie blue. The size of the gel piece containing bands to be cut out from the gel depends on the density (*37*). Before enzymatic digestion of the proteins, in-gel reduction and alkylation reactions are carried out to prevent oxidation and disulfide formation. Subsequently, trypsin was applied for protein digestion. Following enzymatic digestion the samples are transferred to lobind tubes (Eppendorf) before mass-spectrometry analysis as the use of these tubes significantly reduces the binding of hydrophobic proteins to the wall of the tubes (*37*). After enzymatic reaction, all samples were desalted using Sep-Pak C18 cartridges.

### Mass Spectrometry

Peptide fractions were analyzed on an Orbitrap Fusion Lumos Tribrid Mass Spectrometer (Thermo Fisher Scientific) interfaced with Easy-nLC 1200 nanoflow LC system (Thermo Fisher Scientific). The peptides were reconstituted in 0.1% formic acid and loaded on an analytical column at a flow rate of 300 nL/min using a linear gradient of 10-35% solvent B (0.1% formic acid in 95% acetonitrile) over 120 min. Specific conditions for mass spectrometry were set as following; MS1 resolution (120,000), MS2 resolution (30,000), fragmentation method (HCD), collision energy for MS2 (32%). The fifteen most intense precursor ions from a survey scan were selected for MS/MS fragmentation using higher energy collisional dissociation (HCD) fragmentation with 32% normalized collision energy and detected at a mass resolution of 30,000 at 400 m/z. Automatic gain control for full MS was set to 1 × 10^6^ for MS and 5 × 10^4^ ions for MS/MS with a maximum ion injection time of 100 ms. Dynamic exclusion was set to 30 sec. and singly charged ions were rejected (*38*).

### Polysome profiling

Fly brain polysome profile was performed as described by Darnel et al. with minor changes (*39*). Brain tissue homogenates of α-Syn transgenic and age matched control flies (5 days old) were examined and profiled for polysome to understand the abundance translating mRNAs in the polysome fractions. Fly brains were lysed in polysome lysis buffer (10 mM Tris pH 7.5, 150 mM NaCl, 5 mM MgCl_2_, 0.5 mM DTT, 100 mg/ml cycloheximide, 40 U/ml RNase inhibitor, 0.1% sodium deoxycholate and complete protease inhibitor, ROCHE). Approximately ∼ 100 fly heads were lysed in 1 ml polysome lysis buffer, protein concentrations of the lysates were measured and 0.5 mg equivalent polysome lysates were loaded for 10-45% sucrose gradient. The sucrose gradients with polysome lysates were centrifuged at 40,000 rpm for 2 hours at 4°C. All gradients were fractionated in polysome Gradient Station (BIOCOMP) under same parameters all samples. Polysome profiles of HEK293 cells were performed using same buffer and assay conditions. Transfected cells (with 1 μg of DNA per well in six well plate) were harvested after 24 hours and lysed in polysome lysis buffer. Each polysome assay was performed using 0.5 mg total protein.

### Drosophila stocks and maintenance

Drosophila stocks were maintained according to the standard laboratory protocols. The fly stocks were maintained in standard Drosophila diet containing yeast extract, agar, cornmeal, sucrose, and dextrose. For routine maintenance flies were kept in a 12 h light/dark cycle and all fly stocks and experiential crosses were housed at 25°C. UAS α-Syn, UAS A53T-α-Syn and relevant Gal4 stocks were obtained from Bloomington Drosophila Stock Center (BDSC), Indiana, USA. QUAS-α-Syn transgenic fly was a gift from Dr. Mel Feany, Harvard Medical School. Human eIF4E-BP1 and TSC2 constructs were generated in the lab and subcloned the cDNA sequences of eIF4E-BP1 and TSC2 into pQUAST-att plasmid. The verified constructs were used for microinjection into w^118^ embryos (The BestGene, Inc). For experimental control, the relevant Syb-QF2/+ and Gal4/+ (heterozygous) flies were used as controls in the experiments.

### Drosophila climbing and life span assay

Climbing assay was performed as previously described (*40*). Both test and control flies were collected immediately after eclosion and maintained in regular fly food vial at 25°C fly incubator with recommended humidity and aeration. Approximately ∼25 flies were housed in each fly vials. Climbing assays were performed every 3 days by transferring 25 flies to an empty 9 cm fly vial (Genesee Scientific, cat no. 32–113) with a line drawn 6 cm from the bottom. To induce and test an innate climbing response, flies were tapped to the bottom. The number of flies were counted that crossed the 6 cm mark in 15 sec time. For each batch five to ten technical replicates were performed to ensure an accurate reading at each time-point and average of the independent trials was used to calculate the percentage of flies with climbing defects. For fly lifespan assay Syb-QF2>α-Syn flies were collected immediately after eclosion and maintained in the fly food vials with drugs of interest and their vehicles at 25°C. The flies were aged in a batch of 30 per vial and total 120 flies were aged for each drug and corresponding vehicles. In every three days flies were transferred to fresh fly food with drug and surviving flies were counted every five days. Percent fly survival was plotted using Kaplan Meyer survival curve in Graph Pad Prism 9 software.

### Fly brain immunostaining

Fly brains were dissected and TH immunostained following previously described protocols with minor modifications (*41*). Adult Drosophila brains were dissected in PBS and fixed in 4% paraformaldehyde (PFA) at room temperature (RT) for 30 mins. After incubation with PFA, the brains were washed for 30 mins in 1X PBS with 0.3% Triton X-100 (wash buffer), blocked for 30 mins in blocking buffer (1X PBS, 0.3% Triton X-100 and 5% normal goat serum) at RT. Following blocking step, the brains were incubated with primary antibody in blocking buffer for 48 hours at 4°C. The brains were washed three times, 10 mins each, at RT. Secondary antibody in blocking buffer was added to the brains and incubated for 24 hours at 4°C. After five 10 mins washes, brains were carefully mounted on glass slides using ProLong Gold antifade mountant (Life Technologies). The TH (anti-Tyrosine Hydroxylase) primary antibody was used in 1:500 dilution (Immunostar, cat no. 22941), Secondary antibody goat anti-mouse was used at 1:1000 (Thermo Fisher, cat no. A-11032).

### Immunostaining of primary cortical neurons

Primary cortical neurons on glass coverslip in 12-well plate were treated with 5 μg/mL α-Syn PFF in neural basal media for 7 days. The cells on the coverslip were washed with 1X PBS to remove residual media and fixed with 4% paraformaldehyde (PFA) for 15 mins and incubated at room temperature (RT). Following PFA treatment cells were washed with 1X PBS for three 10 mins washes and blocked in 1X PBS with 5% normal goat serum and 0.3% Triton X-100 (blocking buffer) at RT. Primary antibodies were added in blocking buffer for 24 hours at 4°C. The coverslips were washed in 1X PBS with 0.3% Triton X-100 (Wash buffer) for three 10 mins washes at RT. Secondary antibodies were added to the coverslips in blocking buffer and incubated for 1 hour at room temperature followed by the 5 ten mins washes. The coverslips were mounted on glass slides with mounting solutions. The rabbit anti-α-Syn (phospho S129) antibody (Abcam) and mouse anti-MAP2 (Millipore) primary antibodies were used in 1:500 dilution (Millipore. 22941), Secondary antibody goat anti-mouse and anti-rabbit antibodies were used at 1:1000 (Thermo Fisher, cat no. A-11032).

### Toxicity test in primary cortical neuron

Mature primary cortical neurons were treated with 5 μg/mL α-Syn PFF in neural basal media for 14 days. Cell death was measured by treating the neurons with 7 mM Hoechst 33342 and 2 mM propidium iodide (PI) (Invitrogen). Images were captured by a Zeiss microscope and percent cell death was counted using ImageJ software.

### Rapamycin, S6K Inhibitor and Anisomycin treatment for *Drosophila*

The drugs were given to the experimental flies in standard fly food. The flies were treated in batches of 20-25 per fly tube containing 0.5 μM rapamycin (LC laboratories), 10 μM S6K inhibitor (PF-4708671; Pfizer) and 10 μM anisomycin (Sigma) respectively. The drugs were added, in a 200 μL volume, to the dry surface of fly media and allowed to absorb and air dry at least 6 hours prior to the experimental treatment. For both climbing and survival assays the flies were treated for 3 days in a tube and then transferred to tubes containing fresh food and drugs.

### Tissue homogenization and Western blotting

Brain tissues, both human post mortem PD brains as well as mouse brains, were homogenized in 1X RIPA buffer (50 mM Tris HCl, 150 mM NaCl, 1.0% (v/v) NP-40, 0.5% (w/v) Sodium Deoxycholate, 1.0 mM EDTA, 0.1% (w/v) SDS and 0.01% (w/v) sodium azide at a pH of 7.4.) with additional phosphatase inhibitor mixture I and II (Sigma-Aldrich, St. Louis, MO), and complete protease inhibitor mixture (Roche, Indianapolis, IN.). Following homogenization, the samples were freeze-thawed three times using dry ice, centrifuged 15000 x g for 15 min and collected clear supernatants. Protein concentrations were measured using the BCA assay (Pierce, Rockford, IL), SDS-PAGE was used to separate proteins and transfer to nitrocellulose membranes for immunoblot analysis. For blocking, membranes were incubated in 5% non-fat milk in TBS-T (Tris-buffered saline with 0.1% Tween-20) at least 1 h at room temperature (RT) and subsequently probed using primary antibodies overnight at 4°C with continuous shacking. After overnight primary antibody. The membranes were washed 30 mins ---appropriate HRP-conjugated secondary antibodies (Cell signaling, Danvers, MA). The bands were visualized by ECL substrate.

### Isolation of A53T α-Syn aggregates from transgenic mice brain tissues

Freshly dissected brain stem tissues (∼30 mg) from late symptomatic (13 months old) A53T α-Syn transgenic and age matched control mice were homogenized in lysis buffer (10 mM Tris pH 7.5, 150 mM NaCl, 5 mM MgCl_2_, 0.5 mM DTT, 100 μg/ml cycloheximide, 40 U/ml RNase inhibitor, 0.1% sodium deoxycholate and complete protease inhibitor from ROCHE). Tissue lysate volume was adjusted to 600μL for ultracentrifugation at 70K rpm in Optima Max Ultracentrifuge (Bechman Coulter). For ultracentrifugation 45% sucrose solution was prepared in the same lysis buffer. The tissue homogenates were ultracentrifuged in Polycarbonate Thick Walled 13 X 56 mm centrifuge tube (Beckman Coulter), containing 0.9 ml 45% sucrose solution and 0.6 ml tissue homogenate on top of sucrose solution. Centrifugation was performed for 2 hours at 4°C. After centrifugation supernatant was decanted and hard-shiny pellets were resuspended in 100 μL lysis buffer. The preps were dialyzed using MINI Dialysis Device, 10K MWCO (Thermo) in 1L PBS overnight. After overnight dialysis the dialysis device was transferred to fresh 1L PBS to repeat the process overnight. After 48 hours of dialysis the preps were filter sterilized and stored at -80°C.

### Transmission Electron Microscopy (TEM) of Drosophila brains

Drosophila heads were fixed with 4% paraformaldehyde (freshly prepared from EM grade prill) 2% glutaraldehyde 100 mM Sorenson’s phosphate buffer with 5 mM magnesium chloride pH 7.2, overnight at 4 C. Following buffer rinses, samples were microwave fixed twice in 2% osmium tetroxide reduced with 1.5% potassium ferrocyanide, in the same buffer. Sample temperatures did not exceed 9°C. Following microwave processing samples were rocked in osmium on ice for 2 hours in the dark. Tissue was then rinsed in 100 mM maleate buffer with pH 6.2, then en-bloc stained for 1 hour with filtered 2% uranyl acetate in maleate buffer, pH 6.2. Following en-bloc staining samples were dehydrated through a graded series of ethanol to 100%, transferred through propylene oxide, embedded in Eponate 12 (Pella), and cured at 60°C for two days. Sections were cut on a Riechert Ultracut E microtome with a Diatome Diamond knife (45 degree). 60 nm sections were picked up on formvar coated 1 x 2 mm copper slot grids and stained with methanolic uranyl acetate, followed by lead citrate. Grids were viewed on a Hitachi 7600 TEM operating at 80 kV and digital images captured with an XR80-8-megapixel CCD by AMT.

### Antibodies for SiMPull Assay

All antibodies used for SiMPull were obtained from commercial sources as follows: biotinylated anti-RFP from Abcam (ab34771), anti-YFP and anti-Flag from Rockland Antibodies and Assays and anti-myc from Sigma-Aldrich. Primary antibodies for TSC1 and TSC2 were obtained from Cell Signaling Technology. Alexa-647 tagged goat anti-rabbit IgG was purchased from ThermoFisher Scientific. Neutravidin and BSA were procured from Thermo-Fisher and New England Biolabs respectively.

### Cell Culture and Transfection for SiMPull Assay

Assembly of functional mTORC2 was achieved by co-expression of mTOR, Rictor, mSin and mLST8, while mTORC1 and TSC complexes were generated via co-expression of mTOR, Raptor and TSC1, TSC2 respectively. HEK293 cells were grown in DMEM containing 10% (vol/vol) FBS and 2 mM L-glutamate at 37° C with 5% (vol/vol) CO_2_. Transfection of plasmids was carried out using Lipofectamine 2000 (Thermo Fisher Scientific) following the manufacturer’s protocol when the cells reached 60-70% confluence in 6-cm plates. A day after transfection, cells were lysed in 300 μL of lysis buffer containing 40 mM Hepes, pH 7.5, 120 mM NaCl, 10 mM sodium pyrophosphate, 10 mM β-glycerophosphate, 1X protease inhibitor mixture and 0.3% CHAPS. PKA complexes were obtained from HEK cells, transiently transfected with R-Flag-mCherry and C-HA-YFP constructs. PKA-RIIβ and Cα isoforms were used as the regulatory and catalytic subunits respectively. After 24 h expression, cells were lysed using a buffer containing 10mM Tris pH 7.5, 1% NP-40, 150 mM NaCl, 1 mM EDTA, 1 mM benzamidine, 10 μg/ml leupeptin, 1mM NaF, 1mM Na_3_VO_4_. The lysate thus obtained was centrifuged at 14,000 X g for 20 min and subsequently used for SiMPull (*21*).

Pre-determined concentrations of monomers and PFFs of α-synuclein were added to the cell-lysates and incubated at 4° C for 3h. The cell lysates were diluted 50-150 fold in T50-BSA buffer (10 mM Tris, pH 8, 50 mM NaCl and 0.2 mg/mL BSA) to obtain a surface density optimal for single-molecule analysis (∼ 600 molecules in 5,000 μm^2^ imaging area) (*20, 21*).

### Single-Molecule Imaging and Analysis

Single-molecule experiments were performed on a prism-type TIRF microscope equipped with an electron-multiplying CCD camera (EM-CCD) (*42*). For single-molecule pull-down experiments quartz slides and glass cover slips were passivated with 5000 MW methoxy poly-(ethylene glycol) (mPEG, Laysan Bio) doped with 2-5 % 5000 MW biotinylated PEG (Laysan Bio). Each passivated slide and cover slip was assembled into flow chambers. The cell lysates were pulled down with biotinylated antibodies against Flag, myc, YFP or RFP, already immobilized on the surface via neutravidin-biotin linkage. TSC1 and TSC2 were visualized using Alexa-647 tagged anti-rabbit secondary antibodies. YFP-, m-Cherry- and Alexa-647 tagged proteins were excited at 488 nm, 568 nm, and 640 nm respectively and the emitted fluorescence signal was collected via band pass filters (HQ 535/30, Chroma Technology for YFP, BL 607/36, Semrock for mCherry and 665LP from Semrock for Cy5). 15 frames were recorded from each of 20 different imaging areas (5,000 μm^2^) and isolated single-molecule peaks were identified by fitting a Gaussian profile to the average intensity from the first ten frames. Mean spot-count per image for YFP and mCherry was obtained by averaging 20 imaging areas using MATLAB scripts. All experiments were carried out at room-temperature (22-25° C).

The complexes identified were next subjected to photobleaching step analysis to determine the stoichiometry of mTOR and Raptor/Rictor in mTORC1 and mTORC2. A single photobleaching step can be characterized by an abrupt drop in fluorescence intensity. Single-molecule fluorescent time traces from individual YFP or mCherry spots were manually scored for the number of photobleaching steps and the stoichiometry of the molecules was assessed subsequently (*20, 21*). All images were collected at a time resolution of 100 ms. Each molecule was arrayed based on the number of photobleaching steps (typically 1-3) or was discarded if no distinct photobleaching step could be identified. All spots with no fluorescent signal from either of the fluorophores were rejected. A minimum of 500 molecules acquired from at least four different imaging areas were analyzed for each experimental condition.

### Rapamycin treatment of α-Syn PFF injected mice

This experiment was performed in compliance with the regulations of the Animal Ethical Committee of the Johns Hopkins University Animal Care and Use Committee. Rapamycin was administered to α-syn PFF injected mice model in two different ways. First, to examine α-syn pathology in one month of α-syn PFF injected mouse model, both PBS and α-syn PFF mice were injected intraperitonially with Rapamycin (6mg/Kg; #R-5000, LC laboratories) or vehicle (10% PEG400, 10% Tween 80 in water) for 30 days. Rapamycin initiated at day 1 after PFF injection and continued for 30 days and three times per week. Second, to examine toxicity, α-syn PFF injected mice were fed with rapamycin in mouse chow (5LG6-JL) for six months. Mouse feeding experiment was performed as described previously by Harrison et al. (*32*). Microencapsulated rapamycin (eRAPA) was incorporated into mouse chow and provided by Rapamycin Holdings (San Antonio, TX 78249). Briefly, Rapamycin (LC Labs) was microencapsulated by Southwest Research Institute (San Antonio, Texas) using enteric coating material Eudragit S100 (Ro hm Pharma). Microencapsulation protects rapamycin from digestion in the stomach and the encapsulated rapamycin was administered at 14 mg per kg food (2.24 mg of rapamycin per kg body weight per day). The diet containing only the coating material (Eudragit) used as control food in the experiment.

### Stereotactic Injection of α-Syn PFF

Mice were anesthetized with ketamine (100 mg/kg) and xylazine (10 mg/kg). PBS or α-syn PFF (10 μg/mouse) was injected into the striatum (anteroposterior (AP) = +0.2 mm, mediolateral (ML) = + 2.0 mm, dorsoventral (DV) = +2.6 mm, relative to bregma). A 2 μl syringe was used for injections at a rate of 0.4 μl/min, and post-surgical care was provided after surgery.

### Immunohistochemistry

Immunohistochemistry was performed on 40 mm thick serial brain sections. For α-syn phosphorylation detection, sections were blocked with PBS containing 10% normal goat serum and 0.2% Triton X-100 for 1 hour, and then incubated with primary antibodies to p-α-syn, and MAP2 or Tyrosine hydroxylase (TH) overnight at 4°C, followed by incubations with appropriate fluorescent secondary antibodies conjugated to Alexa-fluor 488, 594 or/and 647. P-α-syn pathology was displayed by tracing all visible immunoreactive inclusions/cells and neurites using Keyence Microscope at 10 × magnification. Fluorescent images with higher magnification (20 × or 40 ×) were acquired by confocal scanning microscopy (LSM880, Carl Zeiss). Images were processed using Zen software (Carl Zeiss), and the signal intensity was measured using ImageJ. For histological studies, free-floating sections were rinsed in Tris-buffered saline (TBS, pH 7.4) and incubated with 0.5% H_2_O_2_ in TBS to inhibit endogenous peroxidase. After blocking in TBS containing 10% goat serum and 0.2% Triton X-100, sections were incubated with Rabbit anti-TH antibody overnight at 4°C, followed by incubation with biotin-conjugated anti-rabbit antibody. After three washes, ABC reagent (Vector laboratories) was added, and the sections were developed using DAB peroxidase substrate (Sigma). Sections were counterstained with Nissl (0.09% thionin). For the quantification, both TH and Nissl-positive DA neurons from the substantia nigra pars compacta (SNpc) region were counted using a computer-assisted image analysis system consisting of an Axiophot photomicroscope (Carl Zeiss) equipped with a computer controlled motorized stage (Ludl Electronics), a Hitachi HV C20 camera, and Stereo Investigator software (MicroBright-Field).

### Behavior Tests

#### Grip Strength

Neuromuscular strength was measured by maximum holding force generated by the mice (Biosed). Animals were allowed to grasp a metal grid with either by their fore and/or hind limbs or both. The tail was gently pulled, and the maximum holding force recorded by the force transducer when the mice released their grasp on the grid. The peak holding strength was digitally recorded and displayed as force in grams.

#### Pole Test

The pole test was used to measure bradykinesia (Karl et al., 2003). A metal rod (2.5 foot long with a 9 mm diameter) wrapped with bandage gauze was used as the pole. The pole test protocol was performed as described previously (Kam et al., 2018): two consecutive days of training consisted of three test trials, followed by the actual test. Each animal was placed directly under the top of the pole (3 inch from the top of the pole) with the head held upwards. Results were expressed in turn time and total time. The maximal cutoff of time to stop the test was 60 s.

#### List of Supplementary Materials

Fig. S1: Monomeric α-syn represses mRNA translation.

Fig. S2: Pathogenic α-syn dysregulates mTORC1 pathway in PD models

Fig. S3: Rapamycin suppresses pathology in α-syn PFF mouse model.

Fig. S4: S6K inhibitor recues α-syn pathology and neurotoxicity in PD models. (

Fig. S5: TSC1-TSC2 complex is stable in presence of increasing concentrations of α-syn monomer.

Fig. S6: Translation repressor eIF4E-BP1 overexpression rescues PD pathology.

Table S1. Biological Process Enriched in A53T α-Syn-APEX MS in HEK293.

Table S2. Biological Process Enriched in common interacting proteins

Table S3: List of key resources or reagents

Movie S1. Climbing assay of *Syb-QF2* >α-Syn files fed with Rapamycin in regular fly food

Movie S2. Climbing assay of control flies (*Syb-QF2*/+) fed with Rapamycin in regular fly food

Movie S3. Climbing assay of *Syb-QF2* >α-Syn files fed with anisomycin in regular fly food.

Movie S4. Climbing assay of control flies (*Syb-QF2*/+) fed with anisomycin in regular fly food.

Movie S5. Climbing assay of *Syb-QF2* >α-Syn files fed with S6K inhibitor in regular fly food

Movie S6. Climbing assay of control flies (*Syb-QF2*/+) fed with S6K inhibitor in regular fly food

Movie S7. Climbing assay of *Syb-QF2* >α-Syn, Syb-QF2 >TSC2 and *Syb-QF2* >α-Syn: *Syb-QF2* >TSC2 flies.

Movie S8. Climbing assay of *Syb-QF2* > α-Syn, *Syb-QF2* > 4EBP1 and *Syb-QF2* >α-Syn: *Syb-QF2* >4EBP1 flies

Data file S1. Two-fold enriched interacting proteins in A53T α-Syn-APEX MS in HEK293.

Data file S2. Two-fold enriched interacting proteins α-Syn PFF MS.

## Acknowledgements

We thank Dr. Mel B. Feany (Harvard Medical School) for providing QUAS-SNCA flies and Dr. Christopher J Potter (Johns Hopkins Medicine) for Syb-QF2 driver flies and pQUAST-att plasmid. The authors thank Noelle Burgess and I-Hsun Wu for creating and assisting with illustrations.

## Authors contributions

Conceptualization, M.R.K, T.M.D. and V.L.D.; Methodology, M.R.K. and X.Y.; Investigation, M.R.K., X.Y., S.-U.K., J.M., S.B., S.S.K., H.W., Y.K. A.J., H.H.K., H.G.; Validation, M.R.K., X.Y.; Formal Analysis, M.R.K., X.Y., S.-U.K., J.M.; Resources, T.H., J.R.-O., J.T., V.L.D. T.M.D.; Writing-Original Draft, M.R.K, T.M.D., V.L.D.; Writing-Reviewing & Editing, M.R.K, T.M.D., V.L.D.; Visualization, M.R.K., X.Y.; Funding Acquisition, T.H., V.L.D. and T.M.D.

## Data and materials availability

### Lead Contact

Further information and requests for resources should be directed to and will be fulfilled by the Lead Contact, Ted M. Dawson, M.D., Ph.D.

### Materials Availability

All biological resources, antibodies, cell lines and model organisms and tools are either available through commercial sources or the corresponding authors. Further information and requests for resources and reagents listed in Table S3 that are not commercially available should be directed to the Lead Contact.

### Data and Code Availability

- The data sets generated during the current study are available within the paper or from the corresponding author on reasonable request.
- This paper does not report original code.
- Any additional information required to reanalyze the data reported in this paper is available from the lead contact upon request.

## Supplementary Materials

**Fig. S1:**
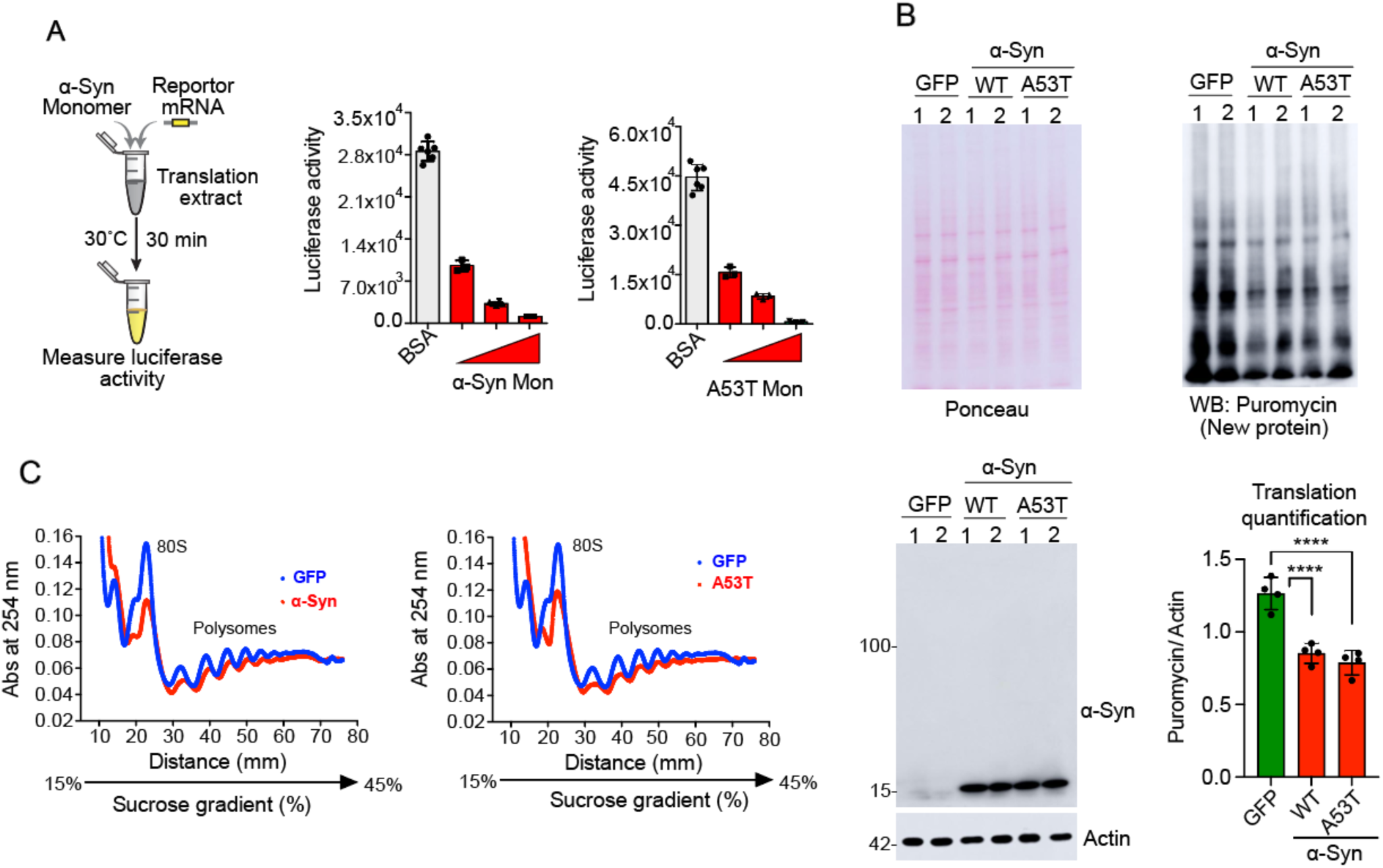
Monomeric α-syn represses mRNA translation. (**A**) Schematic shows experimental design of reconstituted cell free In Vitro Translation (IVT) assay with α-syn monomer. The graph shows readout of IVT reaction incubated with recombinant WT and A53T α-syn monomer with increasing concentrations (n=6). (**B**) SUnSET assay shows overexpression of WT and A53T α-syn in HEK293 cells results in translation repression. The blots show ponceau staining, puromycin and α-syn western in the same blot. The bar graph shows quantification of normalized new protein synthesis (n=4). (**C**) Polysome profiles show association of mRNA with polysome in GFP, WT and A53T α-syn transfected HEK293 cell homogenates. Data are expressed as mean ± SEM; ns, not significant and *p ≤ 0.05, **p ≤ 0.01, and ***p ≤ 0.001. For the figures S2A and S2B statistical significance was measured by One-way ANOVA, data are expressed as mean ± SEM, *p < 0.05, **p < 0.01, ****p < 0.0001.

**Fig. S2:**
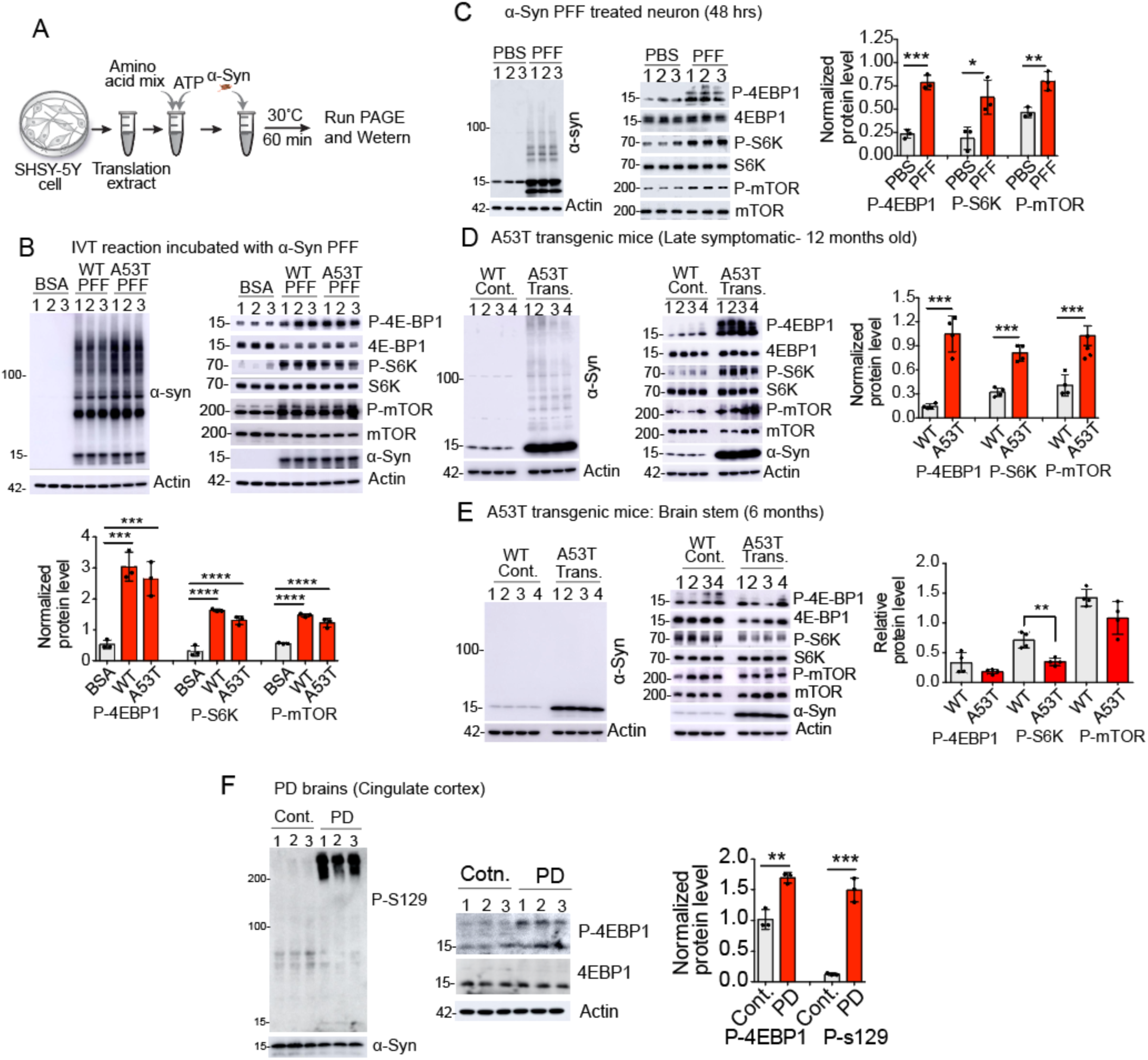
Pathogenic α-syn dysregulates mTORC1 pathway in PD models. (**A**) Schematic depiction of In Vitro Translation (IVT) assay. (**B**) The representative immunoblots show phosphorylated levels of 4E-BP1, S6K and mTOR (S2481). Left blot shows immunoblot of α-syn. The bar graph shows normalized protein levels of P-4EBP1, P-S6K and P-mTOR, n=3. (**C**) Immunoblots show α-syn protein in PFF treated neurons and post translational modifications of mTOR, S6K1 and 4E-BP1 with respective total protein levels. The bar graph shows phosphorylated levels of indicated proteins (n=3). (**D**) Left and middle immunoblot panels show α-syn protein and post translational modification of mTOR, S6K and 4E-BP1 with respective total protein levels. The bar graph shows phosphorylated levels of indicated proteins in the brain stem samples (n=4). (**E**) The Immunoblots show α-syn protein and post translational modification of mTOR, S6K and 4E-BP1 with respective total protein levels in asymptomatic (6 months old) A53T transgenic mice. The graph shows quantification of normalized protein levels as indicated (n=4). (**F**) Immunoblots show eIF4E-BP1 is inactivated in PD brains. Left panel shows protein levels of pathogenic and total α-syn. Right panels show phosphorylated 4E-BP1 in Lewy body rich cingulate cortex from postmortem PD brains. The bar graph shows quantification of pathogenic α-syn and phosphorylated 4E-BP1 protein levels. n=3. For the figures S3C-S3F statistical significance was measured by unpaired two-tailed t test. Data are expressed as mean ± SEM; ns, not significant and *p ≤ 0.05, **p ≤ 0.01, and ***p ≤ 0.001. For the figure S3B statistical significance was measured by One-way ANOVA, data are expressed as mean ± SEM, *p < 0.05, **p < 0.01, ****p < 0.0001.

**Fig. S3:**
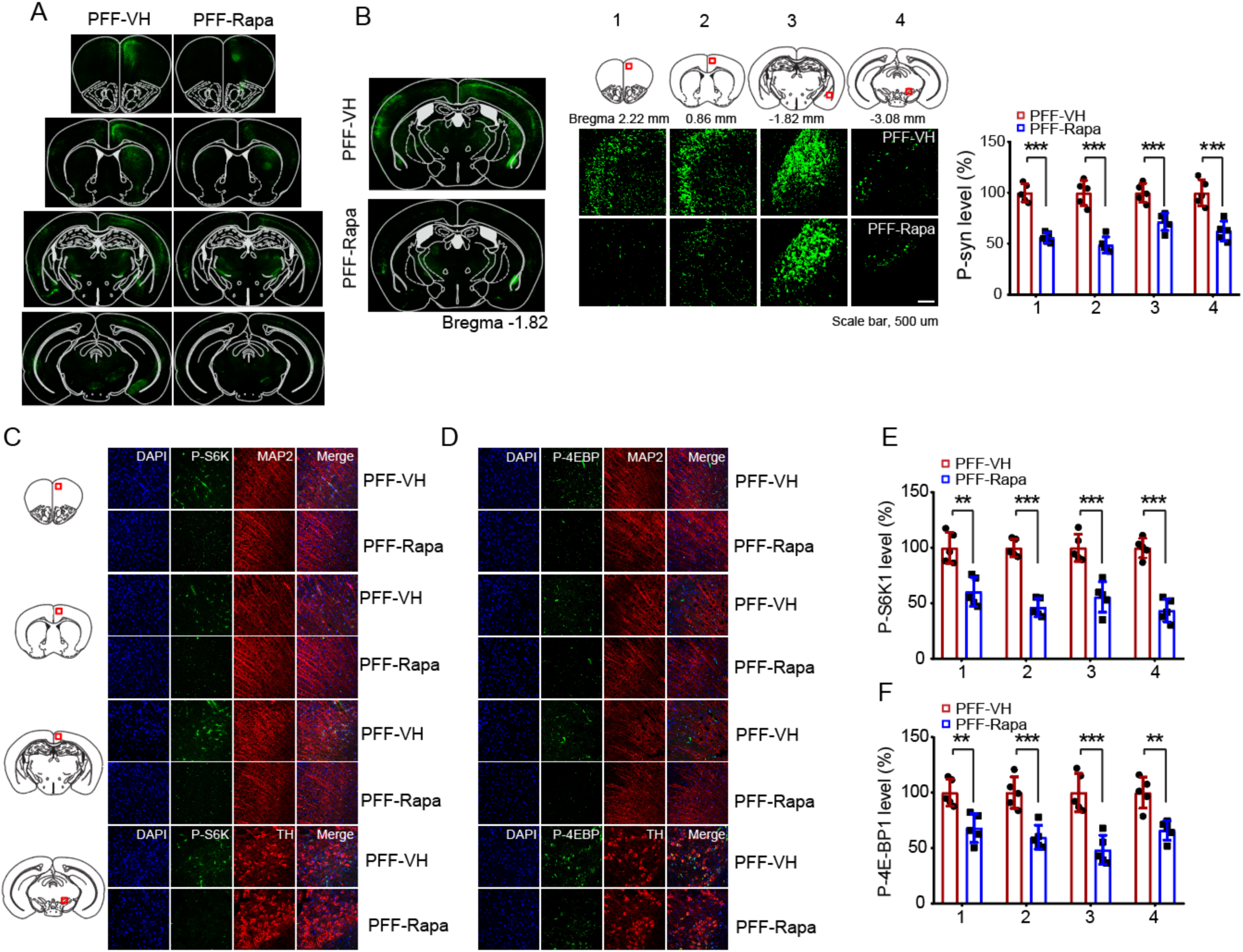
Rapamycin suppresses pathology in α-syn PFF mouse model. (**A**) Whole brain sections of different brain regions of α-syn PFF injected mouse show rapamycin significantly reduces α-syn pathology. (**B**) Zoom-in images show significant reduction of phosphorylated (P-S129) α-syn in the indicated area. The bar graph shows quantification of signal intensity of α-syn pathology (P-S129) (n=5). (**C**) Representative images of P-S6K (P-T389) immunostaining in different brain sections of vehicle and rapamycin injected α-syn PFF mice. (**D**) Immunostaining of P-4EBP1 (P-T37/46) in different brain sections of vehicle and rapamycin injected α-syn PFF mice. **(E)** Quantification P-S6K signal in S Figure 4C (n=5). (**F**) Quantification P-S6K signal in S Figure 4D (n=5). Data are expressed as mean ± SEM; ns, not significant and *p ≤ 0.05, **p ≤ 0.01, and ***p ≤ 0.001.

**Fig. S4:**
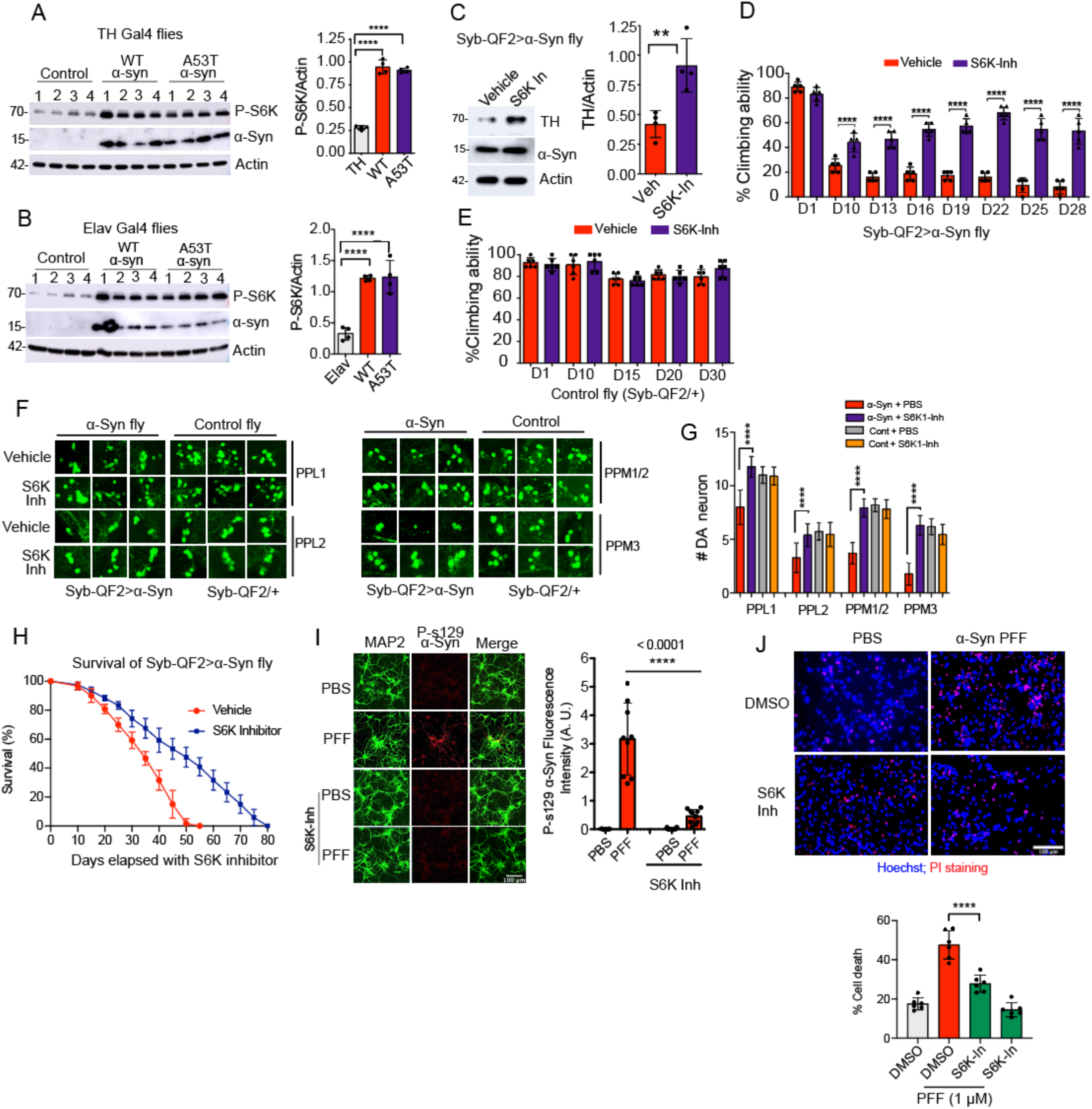
S6K inhibitor recues α-syn pathology and neurotoxicity in PD models. (**A**) Immunoblots shows endogenous *Drosophila* S6K (dS6K) phosphorylation in α-syn transgenic flies (TH Gal4>α-syn and TH Gal4> A53T α-syn compared to the control fly (TH Gal4/+). The bar graph shows quantification of P-dS6K level in fly brain homogenates (n=4). (**B**) Endogenous dS6K phosphorylation in α-syn transgenic flies (Elav Gal4>α-syn and Elav Gal4>A53T α-syn compared to the control flies (Elav Gal4/+). The bar graph shows quantification of P-dS6K levels (n=4). (**C**) Immunoblots show TH, α-syn and actin protein levels in the brain homogenates of S6K-Inh treated fly (Syb-QF2>α-syn). The bar graph shows quantification of normalized TH level (n=3). (**D**) The bar graph shows percent climbing abilities of S6K inhibitor treated α-syn transgenic flies (Syb-QF2>α-syn). Climbing abilities were assessed 28 days. n=5. (**E**) Control flies (Syb-QF2/+) show no adverse effect in motor function due to S6K inhibitor feeding in regular fly food. The graph shows percent climbing abilities of inhibitor treated control flies. Climbing abilities were assessed 30 days. n=5. (**F**) Representative confocal images show individual DA neuron clusters in control (Syb-QF2/+) and α-syn transgenic flies (Syb-QF2>α-syn) after 45 days of S6K inhibitor feeding in regular fly food. (**G**) The bar graph shows the number of DA neurons in the indicated posterior clusters (n=7). (**H**) Survival assay shows increased life span of S6K inhibitor treated Syb-QF2>α-syn transgenic flies (n=4). (**I**) S6K inhibitor prevents pathology in α-syn PFF treated primary cortical neuron. The graph shows quantification of signal intensity of P-S129 α-syn (n=9). Scale = 100 μM. (**J**) Hoechst and PI staining show S6K inhibitor recues toxicity in α-syn PFF treated primary cortical neurons. The bar graph shows quantification of percent cell death (n=6). Scale = 100 μM. For the supplementary figures 5C-5E statistical significance was measured by unpaired two-tailed t test. Data are expressed as mean ± SEM; ns, not significant and *p ≤ 0.05, **p ≤ 0.01, and ***p ≤ 0.001. For the supplementary figures 5A-B, 5G, 5I and 5J statistical significance was measured by One-way ANOVA, data are expressed as mean ± SEM, *p < 0.05, **p < 0.01, ****p < 0.0001.

**Fig. S5:**
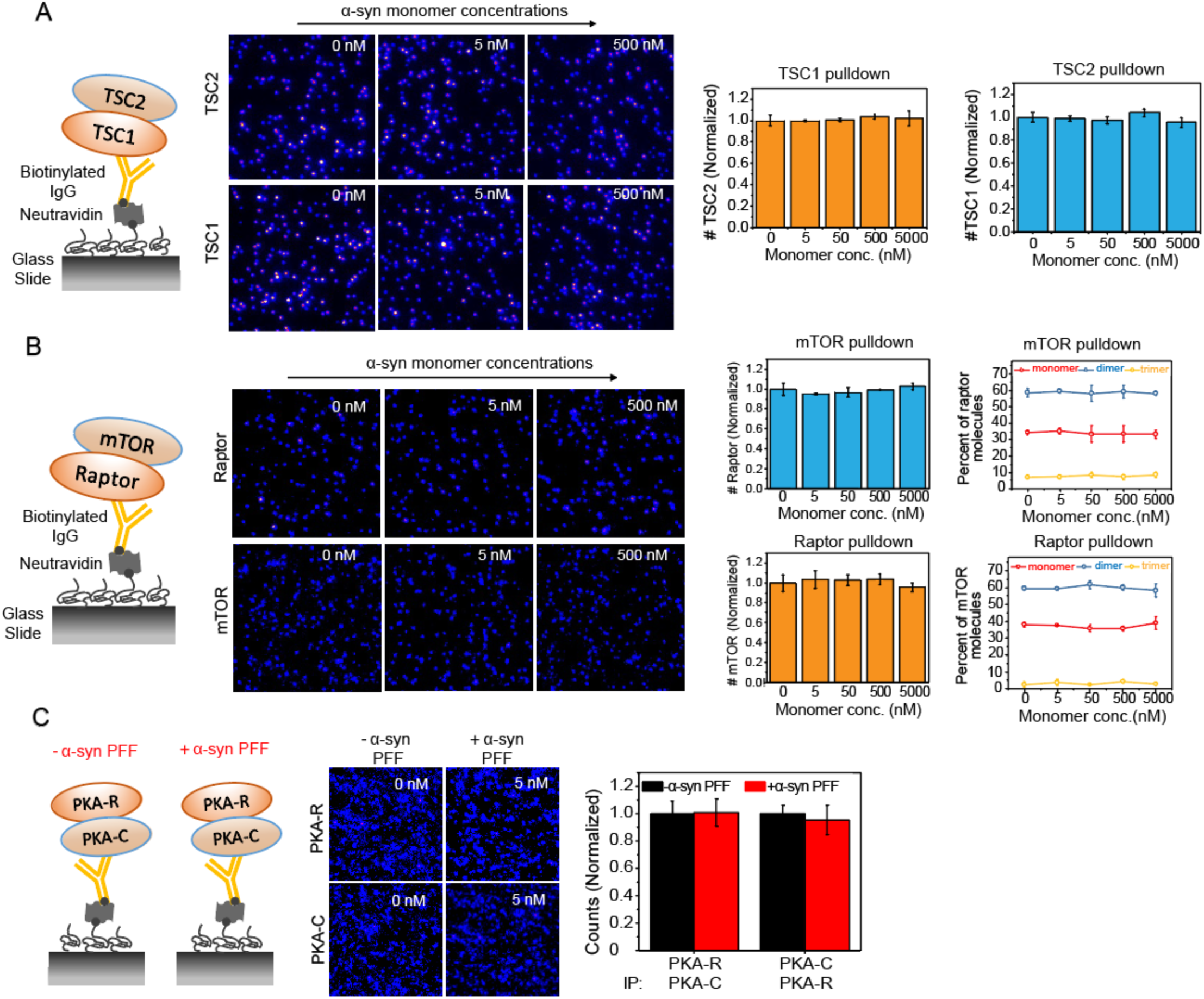
TSC1-TSC2 complex is stable in presence of increasing concentrations of α-syn monomer. (**A**) Schematic shows SiMPull assay design for TSC1-TSC2 complex in presence of α-syn monomer. Representative images show TSC2 and TSC1 molecules pull down by TSC1 and TSC2 respectively in presence of different α-syn monomer concentrations. The bar graphs show normalized levels of TSC2 and TSC1 molecules in SiMPull assay (n=3). (**B**) Schematic shows SiMPull assay design for mTORC1 complex in presence of monomeric α-syn. Representative images show raptor and mTOR molecules pulled down by mTOR and raptor respectively in presence of different α-syn monomer concentrations. The bar graphs show normalized levels of raptor and mTOR molecules (n=3). The line graphs show different species of mTOR and raptor molecules in presence of different α-syn monomer concentrations (n=3). (**C**) Schematic shows SiMPull assay design for PKA complex in presence of pathogenic α-syn PFF. Representative images show molecules of PKA-R and PKA-C. Quantification of PKA-R and PKA-C molecules in the pulldown (n=3). Statistical significance was measured by unpaired two-tailed t test. Data are expressed as mean ± SEM; ns, not significant and *p ≤ 0.05, **p ≤ 0.01, and ***p ≤ 0.001.

**Fig. S6:**
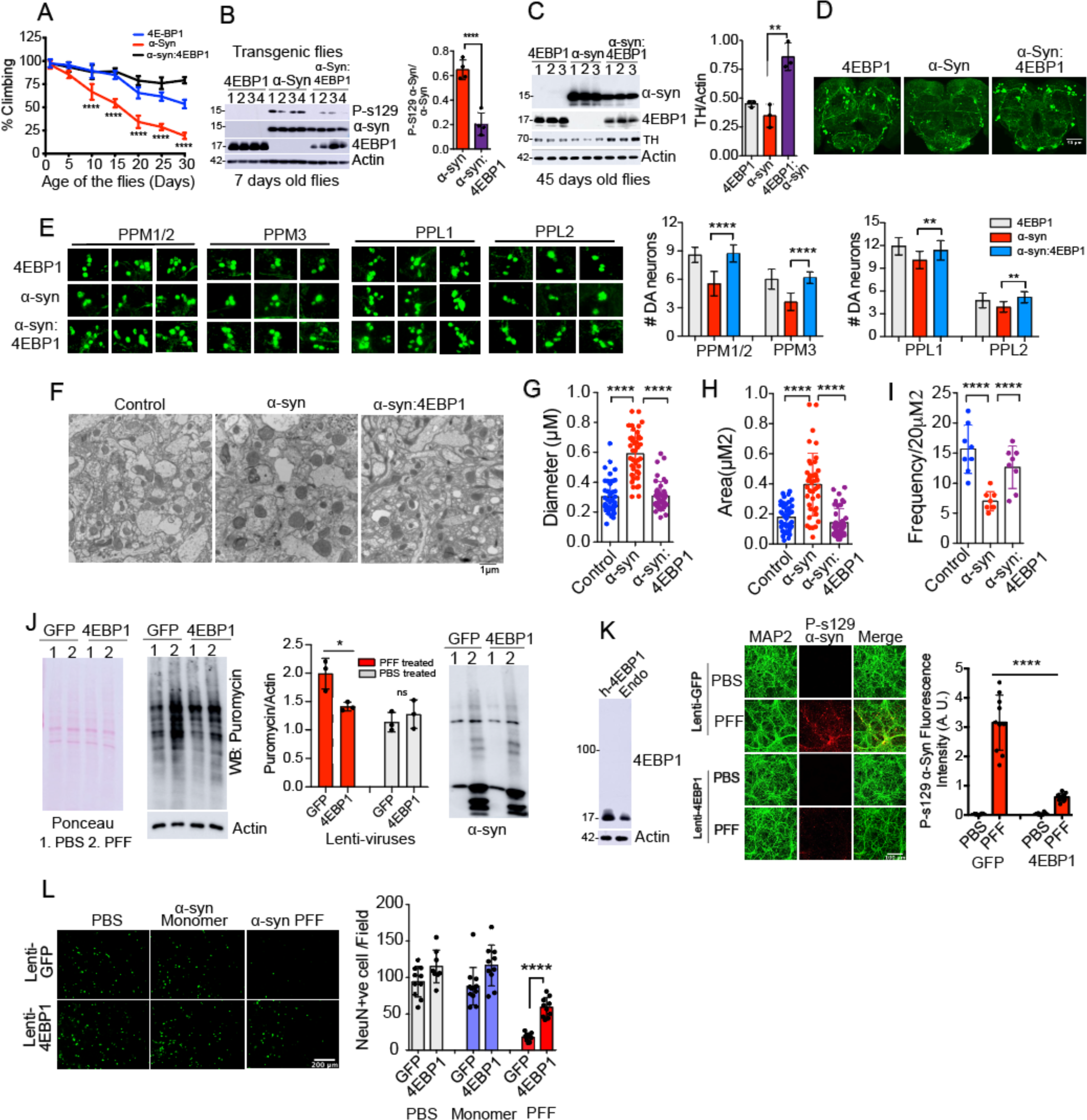
Translation repressor eIF4E-BP1 overexpression rescues PD pathology. (**A**) Line graph shows percent climbing abilities of young and old transgenic flies (Syb-QF2>α-syn; Syb-QF2>4E-BP1 and Syb-QF2>α-syn:4E-BP1). Climbing ability was monitored for 30 days. n=5. (**B**) Overexpression of human eIF4E-BP1 in α-syn transgenic fly shows less pathology. The immunoblots show the protein level of P-s129 α-syn, total α-syn, 4E-BP1 and actin in fly head homogenate. The graph shows quantification of normalized P-s129 α-syn protein level in the immunoblot. n=4. (**C**) Overexpression of 4E-BP1 in α-syn background (Syb-QF2>α-syn:4E-BP1) rescues TH loss. The bar graph shows quantification of normalized TH levels. n= 3. (**D**) Representative confocal images show whole brain TH immunostaining of transgenic flies (Syb-QF2>α-syn:4E-BP1). Scale =50 μM. (**E**) Representative confocal images individual posterior DA neuron clusters in 45 days old single (Syb-QF2>α-syn, Syb-QF2>4E-BP1) and double transgenic (Syb-QF2>α-syn:4E-BP1) flies. The bar graphs present TH positive dopamine neuron counts (n=8). (**F**) TEM micrographs of fly brain cortex compare mitochondrial features (Area, diameter, and frequency) among transgenic flies. Scale = 1 μM. (**G-I**) The bar graphs show quantification of mitochondrial diameter, area, and frequency. (**J**) SUnSET assay shows overexpression of 4E-BP1 in primary cortical neurons suppresses α-syn PFF induced translation. The blots show ponceau staining, puromycin western followed by α-syn and 4E-BP1 immunoblots. The graph shows quantification of puromycin incorporation in newly synthesized protein. n=3. (**K**) The immunoblot shows EIF4EBP1 protein level. The representative confocal images show p-s129 α-syn immunostaining in cortical neuron. The graph shows quantification of signal intensity of P-s129 α-syn (n=9). Scale = 100 μM. (**L**) Overexpression of EIF4EBP1 prevents α-syn PFF induced cell death in mouse primary cortical neuron. Representative confocal images show NeuN staining of live cells. The graph shows quantification of NeuN positive cells (n=10). Scale = 200 μM. For the supplementary figures 6J, 6L and 6M statistical significance was measured by unpaired two-tailed t test. Data are expressed as mean ± SEM; ns, not significant and *p ≤ 0.05, **p ≤ 0.01, and ***p ≤ 0.001. For the supplementary figures 6A-C, 6E, 6G-I and 6K statistical significance was measured by One-way ANOVA, data are expressed as mean ± SEM, *p < 0.05, **p < 0.01, ****p < 0.0001.

## Supplementary Tables

**Table S1.** Biological Process Enriched in A53T α-Syn-APEX MS in HEK293.

| **<u>Biological Process</u>** | **<u>Bonferroni</u>** |
| --- | --- |
| rRNA processing | 1.18E-97 |
| **Translational initiation** | **3.88E-63** |
| Cell-cell adhesion | 2.47E-59 |
| SRP-dependent co-translational protein targeting to membrane | 1.32E-56 |
| mRNA splicing, via spliceosome | 2.19E-54 |
| **Translation** | **8.56E-53** |
| Nuclear-transcribed mRNA catabolic process, nonsense-mediated decay | 2.39E-52 |
| RNA splicing | 1.67E-27 |
| mRNA processing | 1.61E-23 |
| mRNA export from nucleus | 9.11E-22 |
| Mitochondrial translational elongation | 4.33E-20 |
| Regulation of mRNA stability | 2.54E-19 |
| RNA processing | 4.14E-18 |
| Mitochondrial translational termination | 4.99E-18 |
| Protein sumoylation | 1.01E-15 |
| Regulation of cellular response to heat | 2.66E-15 |
| Termination of RNA polymerase II transcription | 2.66E-15 |
| mRNA 3'-end processing | 2.11E-14 |
| RNA secondary structure unwinding | 3.09E-14 |
| RNA export from nucleus | 6.01E-14 |

**Table S2.** Biological Process Enriched in common interacting proteins

| **<u>Biological Process</u>** | **<u>Bonferroni</u>** |
| --- | --- |
| Cell-cell adhesion | 2.81E-20 |
| **tRNA aminoacylation for protein translation** | 1.86E-10 |
| Actin filament organization | 8.93E-07 |
| Positive regulation of canonical Wnt signaling pathway | 9.31E-06 |
| SRP-dependent co-translational protein targeting to membrane | 6.35E-05 |
| Actin filament-based movement | 8.64E-05 |
| Actin cytoskeleton organization | 1.12E-04 |
| **Translational initiation** | **1.67E-04** |
| Endocytosis | 1.86E-04 |
| Nuclear-transcribed mRNA catabolic process, nonsense-mediated decay | 3.29E-04 |
| Viral process | 4.16E-04 |
| Microtubule cytoskeleton organization | 5.52E-04 |
| Actomyosin structure organization | 5.69E-04 |
| Osteoblast differentiation | 7.68E-04 |
| Actin filament bundle assembly | 8.59E-04 |
| Protein phosphorylation | 9.80E-04 |
| rRNA processing | 0.00111015 |
| **Translation** | **0.00113458** |
| Viral transcription | 0.00118906 |
| Negative regulation of translation | 0.00146755 |
| Wnt signaling pathway | 0.00159044 |
| Positive regulation of protein catabolic process | 0.00170897 |

**Table S3:**
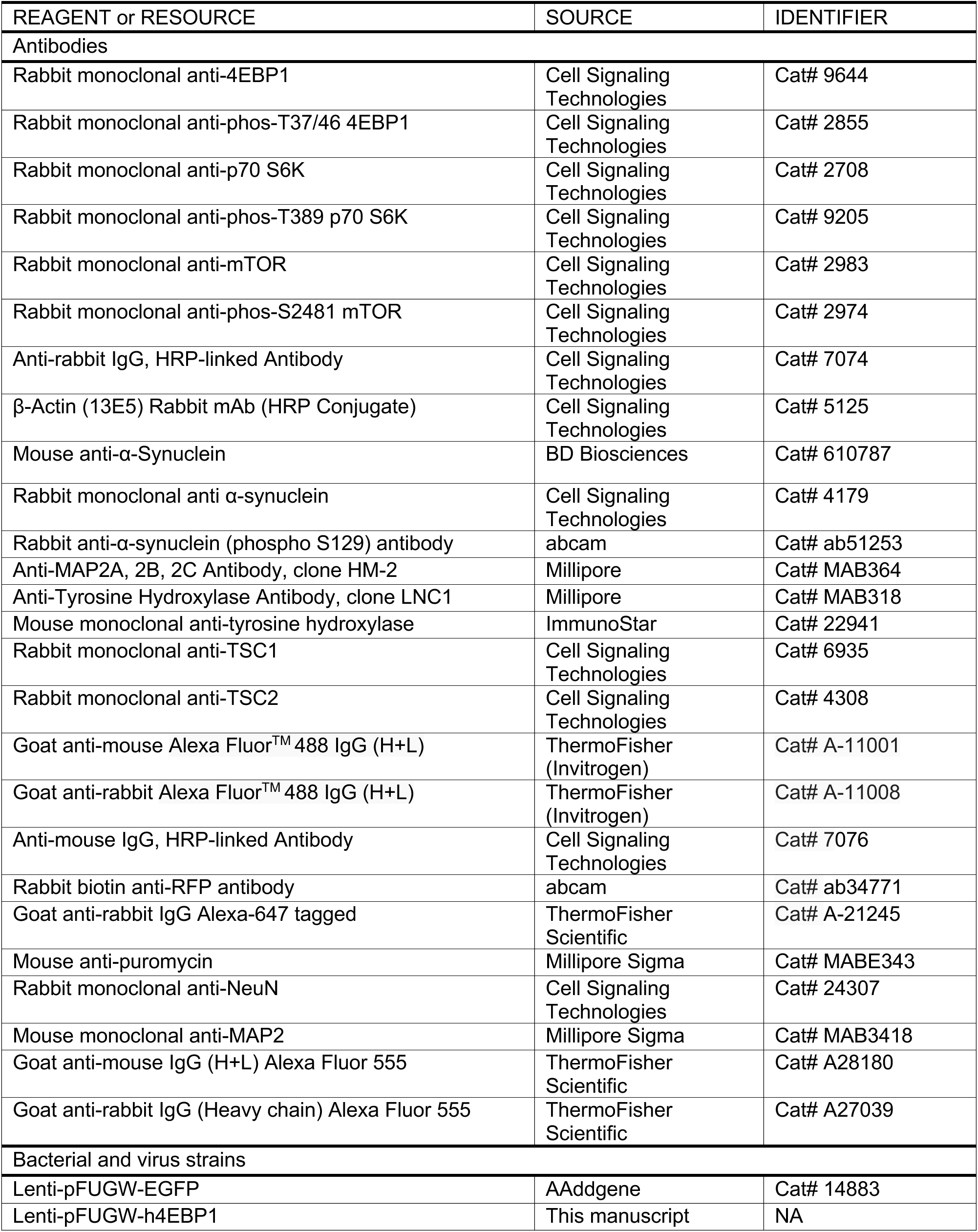
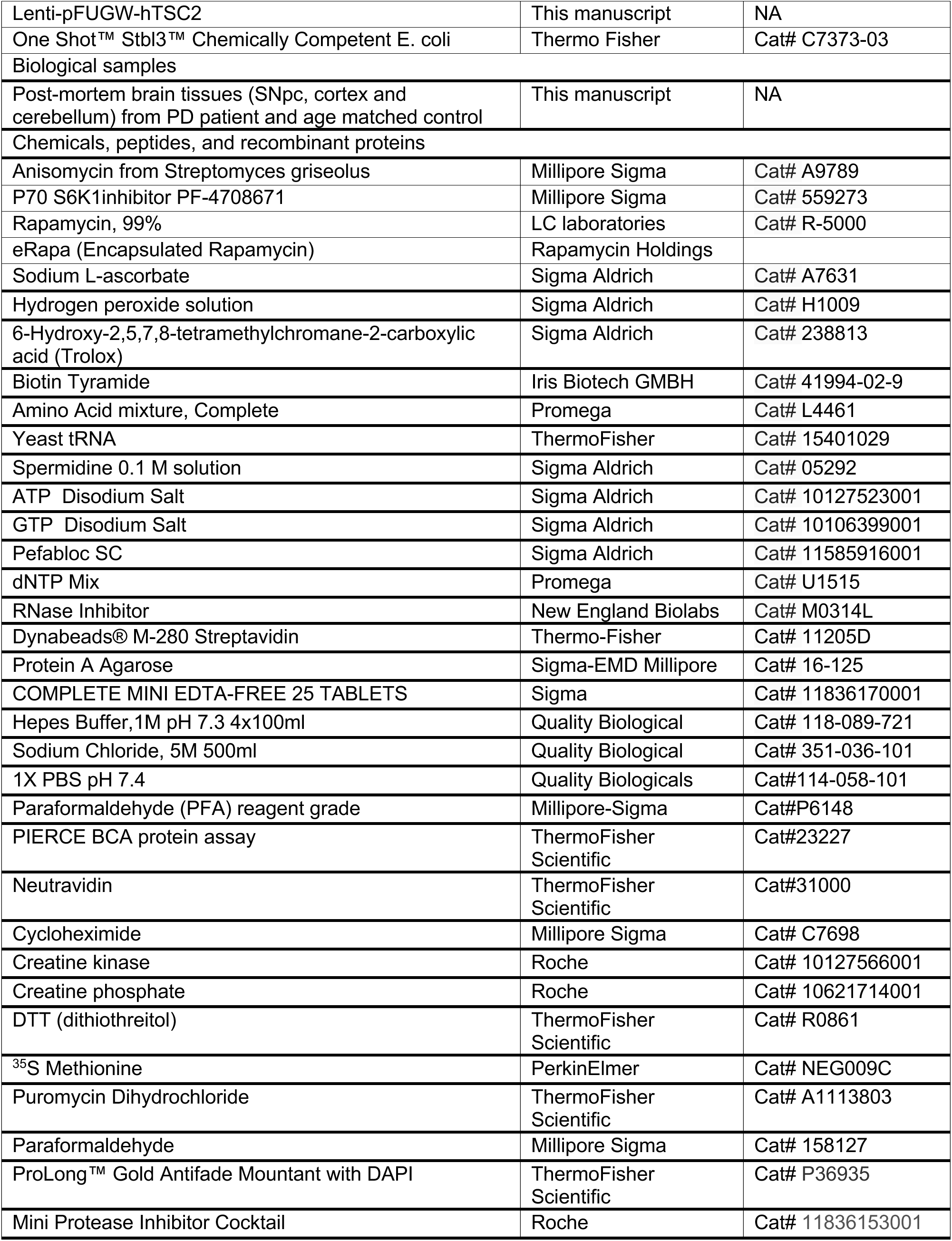
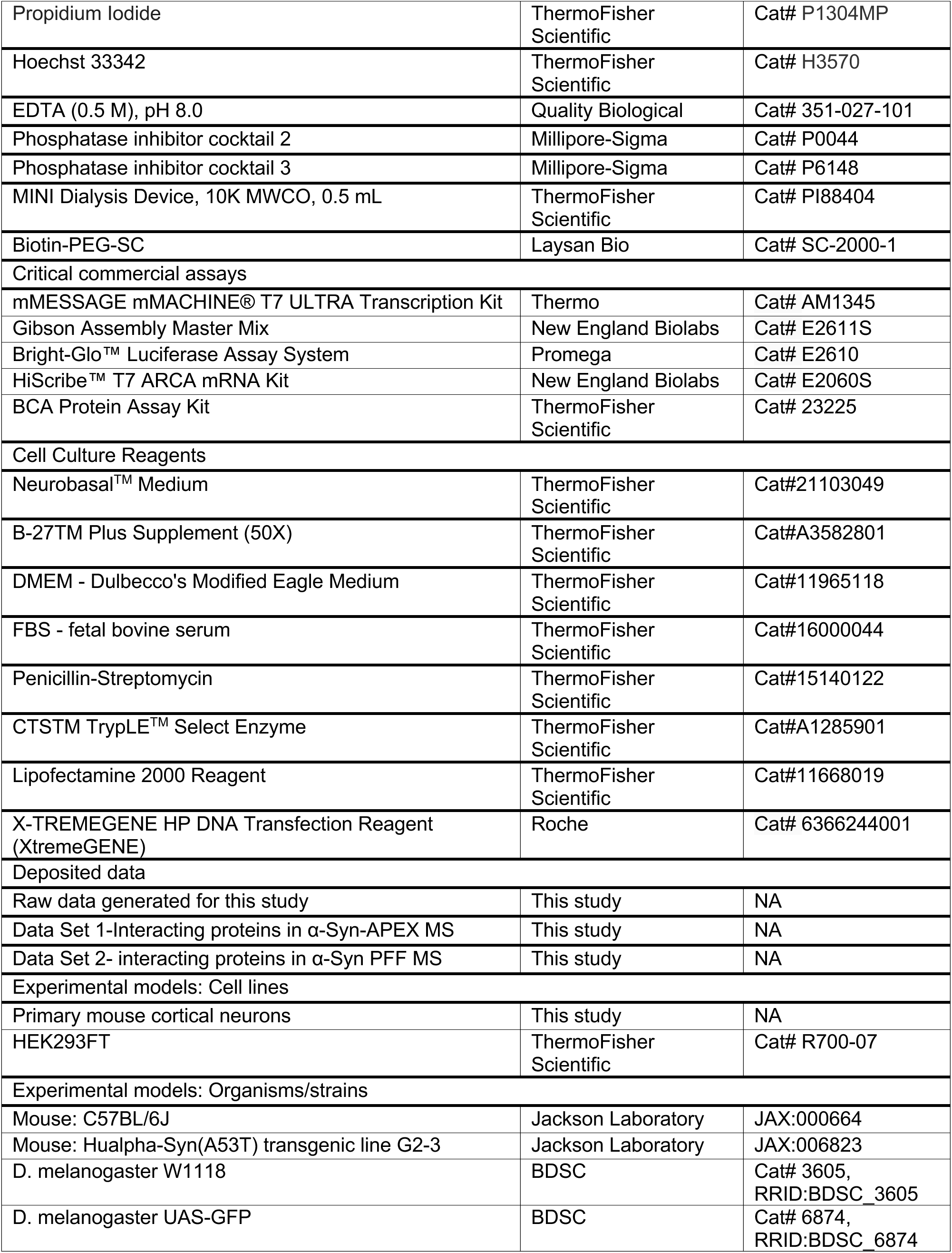
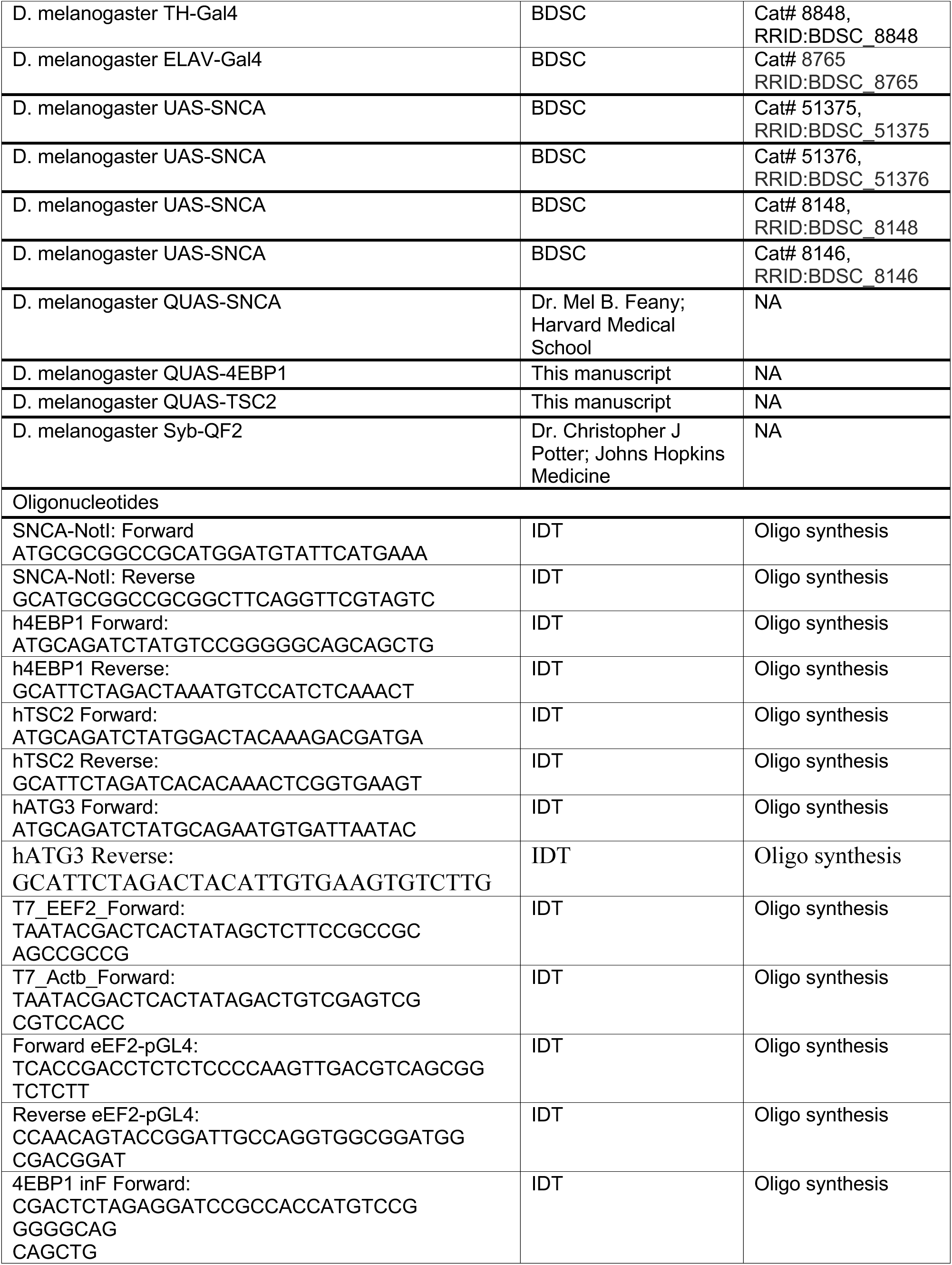
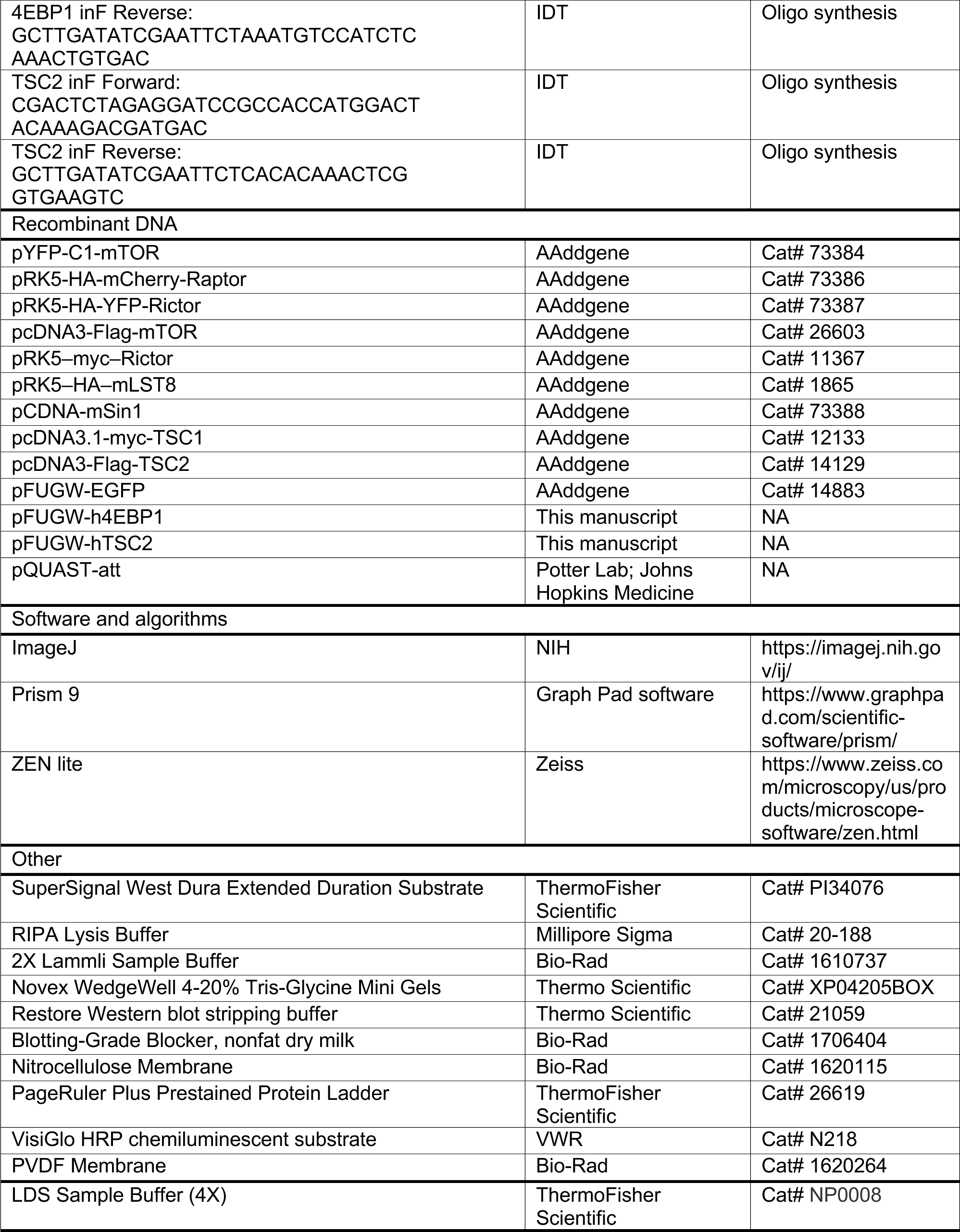
List of key resources or reagents

## Supplementary Movie Legends

Movie S1. Climbing assay of *Syb-QF2* >α-Syn files fed with Rapamycin in regular fly food

Movie S2. Climbing assay of control flies (*Syb-QF2*/+) fed with Rapamycin in regular fly food

Movie S3. Climbing assay of *Syb-QF2* >α-Syn files fed with anisomycin in regular fly food.

Movie S4. Climbing assay of control flies (*Syb-QF2*/+) fed with anisomycin in regular fly food.

Movie S5. Climbing assay of *Syb-QF2* >α-Syn files fed with S6K inhibitor in regular fly food

Movie S6. Climbing assay of control flies (*Syb-QF2*/+) fed with S6K inhibitor in regular fly food

## Supplementary Data Files

**Data file S1. Two-fold enriched interacting proteins in A53T α-Syn-APEX MS in HEK293**

**Data file S2. Two-fold enriched interacting proteins α-Syn PFF MS**

